# Inhibitor of cardiolipin biosynthesis-related enzyme MoGep4 confers broad-spectrum anti-fungal activity

**DOI:** 10.1101/2024.06.16.599186

**Authors:** Peng Sun, Juan Zhao, Gan Sha, Yaru Zhou, Mengfei Zhao, Renjian Li, Xiaojing Kong, Qiping Sun, Yun Li, Ke Li, Ruiqing Bi, Lei Yang, Ziting Qin, Wenzheng Huang, Yin Wang, Jie Gao, Guang Chen, Haifeng Zhang, Muhammad Adnan, Long Yang, Lu Zheng, Xiao-Lin Chen, Guanghui Wang, Toshiki Ishikawa, Qiang Li, Jin-Rong Xu, Guotian Li

## Abstract

Plant pathogens cause devastating diseases, leading to serious losses to agriculture. Mechanistic understanding of pathogenesis of plant pathogens lays the foundation for the development of fungicides for disease control. Mitophagy, a specific form of autophagy, is important for fungal virulence. The role of cardiolipin, mitochondrial signature phospholipid, in mitophagy and pathogenesis is largely unknown in plant pathogenic fungi. The functions of enzymes involved in cardiolipin biosynthesis and relevant inhibitors were assessed using a set of assays, including genetic deletion, plant infection, lipidomics, chemical-protein interaction, chemical inhibition, and field trials. Our results showed that the cardiolipin biosynthesis-related gene *MoGEP4* of the rice blast fungus *Magnaporthe oryzae* regulates growth, conidiation, cardiolipin biosynthesis, and virulence. Mechanistically, MoGep4 regulated mitophagy and Mps1-MAPK phosphorylation, which are required for virulence. Chemical alexidine dihydrochloride (AXD) inhibited the enzyme activity of MoGep4, cardiolipin biosynthesis and mitophagy. Importantly, AXD efficiently inhibited the growth of 10 plant pathogens and controlled rice blast and Fusarium head blight in the field. Our study demonstrated that MoGep4 regulates mitophagy, Mps1 phosphorylation and pathogenesis in *M. oryzae*. In addition, we found that the MoGep4 inhibitor, AXD, displays broad-spectrum antifungal activity and is a promising candidate for fungicide development.

## Introduction

Crop diseases pose serious threats to global food security. Rice blast is one of the most devastating diseases of cultivated rice, eliminating enough grain to feed over 60 million people annually. Rice blast is caused by the filamentous fungus *Magnaporthe oryzae*, which uses specialized infection structures known as appressoria for infection (Hamer *et al*., 1988). As a model and devastating fungus, mechanisms involved in *M. oryzae* pathogenesis have been extensively studied, and multiple biological processes involved in functional appressoria have been identified, one of which is mitophagy, a specific form of autophagy. Mitophagy is involved in timely removal of damaged or an excessive number of mitochondria, important for the maintenance of cellular homeostasis (Mamaev and Zvyagilskaya, 2019). In *M. oryzae*, the mitophagy receptor MoAtg24 is required for fungal virulence (He *et al*., 2013, Kou *et al*., 2019). Additionally, mitochondrial dynamics and morphology-related proteins, which are possibly involved in mitophagy, including MoWhi2, MoAuh1, MoFzo1 and MoDnm1, are crucial for virulence in *M. oryzae* (Sentelle *et al*., 2012, He *et al*., 2013, Shen *et al*., 2017, Kou *et al*., 2019, Meng *et al*., 2022). Whether other components involved in mitophagy are important for virulence in *M. oryzae* has not been investigated.

Phospholipids are major components of biological membranes. The phospholipid, cardiolipin (CL) specifically localized in the inner mitochondrial membrane in eukaryotes, is essential for maintaining mitochondrial functions, including respiration, protein transport and stress responses (Gebert *et al*., 2009). In yeast, the cardiolipin biosynthesis includes *de novo* CL biosynthesis and CL remodeling. Genes *TAM41*, *PGS1*, *GEP4*, and *CRD1* are involved in *de novo* CL biosynthesis, and *CLD1* and *TAZ1* acyltransferase are involved in remodeling CL acyl chains (Ji and Greenberg, 2022). Studies have shown that CL is involved in mitophagy in model organisms (Chu *et al*., 2013, Shen *et al*., 2017), and pathways involved in mitophagy have been studied in yeast *Saccharomyces cerevisiae*, including the mitogen-activated protein kinase (MAPK) signaling pathway (Mao *et al*., 2011, Mamaev and Zvyagilskaya, 2019). However, the role of CL and its metabolic enzymes in plant pathogenic fungi remains unknown. In *M. oryzae*, both mitophagy and MAPK signaling pathways are important for pathogenesis, but whether CL and its metabolic enzymes are involved in these pathways and subsequently *M. oryzae* virulence remains elusive.

Here, our results showed that knocking out the cardiolipin biosynthesis-related gene *MoGEP4* resulted in defects in growth, conidiation, CL biosynthesis, and virulence in *M. oryzae*. Furthermore, we show that the *Mogep4* mutants are impaired in mitophagy and Mps1-MAPK activation. Importantly, we have identified a broad-spectrum fungicide candidate, alexidine dihydrochloride (AXD), which inhibited MoGep4 enzyme activity, fungal development and plant infection. Finally, we show that AXD is efficacious in control of rice blast and Fusarium head blight in the field.

## Materials and methods

### Strains and culture conditions

The wild-type (WT) *M. oryzae* P131 strain and transformants were routinely cultured on complete medium (CM) or oatmeal tomato agar medium (OTA) at 28°C (Li *et al*., 2017). All the transformants were selected on TB3 medium containing 250 μg/ml of hygromycin B (Genview, AH169) or 400 μg/ml of geneticin (GLPBIO, GC17427). The strains used in this study are listed in Table S1. Colony diameters were measured at 6 dpi on CM, OTA and TB3 plates. For tolerance assays, colony diameters were measured at 6 dpi on CM plates supplemented with CaCl2 (0.1 M), Calcofluor White (0.1 mg/ml), Congo Red (0.2 mg/ml), ethidium bromide (EtBr, 2.5 µg/ml), H2O2 (0.01 M), NaCl (0.7 M), or sorbitol (1 M). For heat tolerance assays, strains were cultured at 34°C. All the strains were cultured at 28°C in the dark, and the growth rate was measured on complete medium (CM) or potato dextrose agar (PDA) plates supplemented with/without alexidine dihydrochloride (AXD). The pathogens included *Botrytis cinerea* (B05.10), *Bipolaris maydis* (TM17), *Bipolaris sorokiniana* (LanKao9-3), *Fusarium graminearum* (PH-1), *Leptosphaeria biglobosa*, *Monilinia fructicola* (2YTF2-2), *Phytophthora sojae* (P6497), *Valsa mali* (VM-1), and *Valsa pyri* (VA-1).

### Generation of knockout mutants, the point mutation strain and complementation assays

To generate the deletion mutant of *M. oryzae*, a split-marker strategy was performed according to the standard protocol (Catlett *et al*., 2003). Primers used to amplify the fragments for *MoGEP4* deletion are listed in Table S2. The partial coding region of *MoGEP4* with 1.0 kb flanking sequence was used for screening transformants by PCR. To construct the MoGep4-GFP complementation vector, primers Gep4-Com-F/R were used to amplify the *MoGEP4* ORF with its native promoter. The MoGep4-GFP complement vector was transformed into the *Mogep4* mutant protoplasts and selected with the neomycin-resistance marker. Transformants were screened with primers Gep4-5F/pKNT-seqR and further confirmed by microscopic imaging. To generate strains with point mutations in *MoGEP4*, additional primers GEP4-D60A-F/R that introduced point mutations were used to amplify the *MoGEP4* ORF with its native promoter. The construct was then transformed into the *Mogep4* mutant, similar to the complementation assays.

### Conidiation, conidial germination, appressorium formation, and appressorial penetration assays

Conidiation was measured in ten-day-old cultures on CM plates as previously described (Chen *et al*., 2008). For conidial germination and appressorium formation assays, conidia were collected from CM plates and inoculated onto hydrophobic glass coverslips. Percentages of conidial germination and appressorium formation rates were quantified under a microscope (Nikon, 80i).

Appressorial penetration assays were performed on barley epidermal strips as described (Zheng *et al*., 2015). Briefly, fresh conidia were harvested from CM plates, and 10 μl of the conidial suspension (1 × 10^5^ conidia/ml) in 0.025% Tween 20 was inoculated onto 7-day-old barley epidermal strips, which were incubated in a dark, moist chamber at 28°C. Invasive hyphae were observed at 30 hpi under a confocal microscope (Leica TCS SP8).

### Plant infection assays

Three-week-old rice (*Oryza sativa*) seedlings of cultivar Lijiangxintuanheigu (LTH) and 7-day-old barley (*Hordeum vulgare*) seedlings of cultivar E9 were used for infection assays (Liu *et al*., 2018). Drops of conidial suspension (1 × 10^5^ conidia/ml) in 0.025% Tween 20 were inoculated onto leaves in a dark chamber at 28°C and photographed at 5 dpi. The infection assays of *B. cinerea* and *V. pyri* were performed as previously described (Wu *et al*., 2022, Yuan *et al*., 2022). Briefly, conidial suspensions (5 x 10^5^ conidia/ml) of *B. cinerea* and *V. pyri* were inoculated with/without AXD on tomato and pear leaves, respectively. The disease index was recorded at 5 dpi, and the lesion area was statistically analyzed with ImageJ.

### Quantitative reverse transcription-polymerase chain reaction (qRT-PCR)

Mycelia used for qRT-PCR were the same as in the mitophagy analysis. Total RNA was extracted with the RNAprep Pure Plant Kit (TIANGEN, DP432) and used to generate complementary DNA (cDNA) with the 1st Strand cDNA Synthesis Kit (Vazyme, R312-01). qRT-PCR analysis was performed on the CFX Connect Real-time PCR System using TransStart Tip Green qPCR SuperMix (TransGen Biotech, AQ142-11). The data were analyzed by the 2^-ΔΔCt^ method, and the *β*-tubulin gene (MGG_00604) was used as the internal control (Rao *et al*., 2013). Primers are listed in Table S2.

### Complementation of the yeast *gep4* mutant

The yeast strain BY4741 (MATa *his3 leu2 met15 ura3*) was used in yeast mutant complementation assays. Primer pairs Gep4KOF1/R1, Gep4KOF2/R2 were used to amplify the upstream and downstream fragments of *GEP4*, respectively. KanMX amplified from plasmid pmloxp32 was ligated with flanking fragments with the primer pair Gep4SCF1/R1. The yeast endogenous *GEP4* gene was deleted by the lithium acetate/single-stranded carrier DNA/PEG method as previously described (Sweigard *et al*., 1998). The open reading frame (ORF) of *MoGEP4* was amplified from *M. oryzae* complementary DNA (cDNA) and cloned into the pYES2/NTA vector with the CloneExpress II One Step Cloning Kit (Vazyme, C112-01). The resultant pYES2-*GEP4* construct was sequenced and transformed into the yeast *gep4* mutant. The Ura^+^ transformants were verified by PCR. Growth assays were performed on yeast extract-peptone-dextrose (YPD) plates and yeast-peptone-galactose (YPGal) plates at 30°C and 37°C, respectively. Data were collected at 3 and 5 dpi on YPD and YPGal, respectively. Additional primers GEP4-D60A-F/R were used to introduce point mutations into *MoGEP4*, and the yeast complementation assays were performed similarly.

### Lipid extraction and analysis

Fresh *M. oryzae* protoplasts were harvested for isolation of mitochondria. The protoplasts were suspended and washed in the mitochondrial preparation solution (10 mM KCl containing 0.25 M sucrose, 5 mM EDTA, 20 mM HEPES, pH 7.2, and 0.15% bovine serum). Samples were ultrasonically treated under cool conditions for 5 min to disrupt the cells, and then the suspension was centrifuged at 1,500 g for 10 min at 4°C. The collected supernatant was centrifuged at 10,000 g for 20 min, and the supernatant was removed. The precipitate was washed with the mitochondrial preparation solution and resuspended in 1 ml of chloroform/methanol/formic acid (10:10:1, v/v/v). Vortex extraction was conducted in a glass bottle for 5 min, and then the sample was centrifuged at 1,000 g for 6 min at 4°C to collect the supernatant.

Lipid extraction was performed as previously described (Bligh and Dyer, 1959, Sha *et al*., 2023). The precipitate was resuspended in 1 ml of chloroform/methanol/water (5:5:1, v/v/v) extraction solution, and this step was repeated once. The extracted supernatants were added to 1 ml of salt solution (0.2 M H3PO4, 1 M KCl) and the lowest organic phase was collected by votexing and centrifuging. The solvent was evaporated in a N2 gas stream and redissolved in 100 μl of chloroform. Samples were stored at -80°C for later use.

Two-dimensional thin layer chromatography (2D-TLC) was used for lipid analysis. 2D-TLC plates were pretreated at 110°C for 90 min and samples were spotted at one corner of the plates (Macherey Nagel).

Chloroform/ethanol/ammonium hydroxide (65:25:2, v/v/v) and chloroform/ethanol/acetic acid/water (85:15:10:3, v/v/v/v) were used for the first and second dimensional separation, respectively. After drying for 10 min, the plates were exposed to iodine vapor for 90 sec in the chromatography tank.

### Mass spectrometry analysis of lipids

The 10-ng internal CL standard (14:0-14:0-14:0-14:0) (Sigma-Aldrich, 63988-21-6) was added to lipid extracts for lipidomic analyses. LC-MS/MS (multiple-reaction-monitoring mode) analyses were performed with a mass spectrometer QTRAP 4000 (ABSciex) coupled to a liquid chromatography system (LC20A HPLC, Shimadzu) (Ting *et al*., 2019). Analyses were achieved in the positive mode. Lipids were separated on an Accucore C30 (100 × 2.1 mm, particle size, 2.6 μm, Waters) with eluent A and B solutions. Eluent A was water: methanol: acetonitrile:300 mM ammonium acetate = 20:20:20:1 (v/v/v/v), and eluent B was isopropanol: methanol:300 mM ammonium acetate = 180:20:3 (v/v/v). The gradient elution program was as follows: 0-2nd min, 25%-40% eluent B; 2nd-4th min, 40%-95% eluent B; and 4th–18th min, eluent 95% B. The flow rate was set at 0.3 ml/min and the sample volume was 2 μl. The areas of LC peaks were determined with MultiQuant software (ABSciex) for relative quantification.

### Cell wall, glycogen and lipid droplet staining assays

For Calcofluor white (CFW) staining, mycelia were grown in liquid CM for 24 h, stained with 10 µg/ml of CFW (Sigma-Aldrich, 910090) for 10 min (Liu *et al*., 2018), and observed under the epifluorescence microscope (Nikon, 80i).

For glycogen and lipid droplet staining, conidia were inoculated onto hydrophobic glass coverslips and stained with staining solutions (10 mg/ml I2 and 60 mg/ml KI) and 2.5 μg/ml Nile red (50 mM Tris/maleate buffer, pH 7.5, with 20 mg/ml polyvinylpyrrolidone and 2.5 µg/ml Nile Red), respectively (Wang *et al*., 2018). The stained germinated conidia and appressoria were observed under an epifluorescence microscope (Nikon, 80i).

### Mitophagy analysis

To monitor the mitophagy process in *M. oryzae*, the construct of Idh1 fused with the GFP tag was transformed into the WT and *Mogep4* (M60) strains. The MoIdh1-GFP strains were cultured in CM for 48 h, and then the mycelia were transferred into lipid nitrogen starvation (MM-N) medium (He *et al*., 2013). Total proteins were extracted with 1.5 ml of lysis buffer (50 mM Tris, 1 mM EDTA, 150 mM NaCl, 1% Triton X-100, pH 7.4 containing 1 mM PMSF and a protease inhibitor cocktail) from mycelia (200 mg) cultured as described above. Subsequently, samples were separated on 10% SDS-PAGE gels at 60 V for 4–6 h. Proteins were transferred onto a polyvinylidene fluoride (PVDF) membrane at 40 V at 4°C for 3 h. The membrane was used for western blotting as previously described (Zheng *et al*., 2015).

### Phytotoxicity assays

The phytotoxicity assays of AXD were performed on seven-day-old wheat (Huamai 9) and four-week-old rice (CO39) seedlings (Wu *et al*., 2023). AXD was dissolved in DMSO and sprayed evenly onto leaves. The fresh weight of wheat and rice seedlings was determined at 3 dpi. Solvent dimethyl sulfoxide (DMSO) was used as the control.

### Molecular docking

The protein structure of MoGep4 was predicted with AlphaFold2 (David *et al*., 2022). The chemical structure of AXD was retrieved from the PubChem Compound database (https://www.ncbi.nlm.nih.gov/pccompound/). Software Autodock and PyMOL was used for molecular docking as previously described (Park *et al*., 2016), and the final docking conformation with the best affinity was selected for conformation.

### Protein expression and purification

The purification of the Glutathione S-transferase (GST)-tagged protein was performed as described previously (Schäfer *et al*., 2015). Briefly, the full-length *MoGEP4* was cloned into the pGEX-6p-1 vector using a one-step cloning kit (Vazyme, China) and transformed into *E. coli* BL21 (DE3). Expression of the MoGep4-GST fusion protein was induced by the addition of 1-mM isopropyl-β-D-thiogalactopyranoside (IPTG) to Luria Broth (LB) medium and incubated at 18°C for 12 h. The *E. coli* cells were then centrifuged at 10,000 rpm for 15 minutes, and the supernatant was removed. The cell pellets were resuspended and sonicated in phosphate-buffered saline (PBS). After centrifugation at 10,000 rpm for 15 minutes, the protein was purified with GST NUPharose 4B (Nuptec, China). The MoGep4 protein was washed with PBS buffer and eluted with elution buffer (20 mM glutathione, 50 mM Tris-HCl, pH 8.0).

### SPR assays

The Surface plasmon resonance (SPR) assay was performed on the Biacore T200 instrument. MoGep4 was immobilized onto the CM5 sensor chip surface via amine coupling. AXD at concentrations of 1.25, 2.5, 5, 10, and 20 µM were used for SPR assays together with the control.

### nanoDSF assays

Thermal shift assays were performed on the Prometheus NT.48 nano DSF instrument. Sample solutions contained 1 mg/ml protein in PBS buffer and 5% DMSO, and were heated in capillary tubes from 20°C to 95°C at a heating rate of 1°C/min, and fluorescence was recorded at 330 nm and 350 nm.

### MoGep4 enzyme activity assays

The MoGep4 enzyme activity was measured with the EnzChek Phosphatase Assay Kit (Invitrogen). The reaction buffer contained 5 mM Tris, 100 mM DiFMUP, 5 mM Bis Tris, 20 mM sodium acetate, and 5% DMSO. MoGep4 (1 mM) with or without AXD (4 mM) was added to the reaction buffer. The Glutathione S-transferase (GST) protein (1 mM) was used as the control. Measurements were taken every 2 min for 60 min at 358 nm excitation spectrum and 455 nm emission spectrum.

### Field trials

The field trials to determine the effect of AXD on rice blast were performed as previously described in the rice blast nursery at Yangjiang (21°87’ N, 111°98’ E), March 2022 (Liang *et al*., 2021). The four-week-old rice seedlings of the susceptible cultivar CO-39 were transplanted in each 30 m^2^ plot that contained 600 plants 10 cm apart. At 30 d after planting, 6 μM AXD and DMSO (solvent control) were sprayed twice on rice seedlings 4 days apart, and the disease index of rice blast was examined at 16 dpi.

For wheat, the flowering wheat heads of Xiaoyan22 in the field were inoculated with 10 µl of conidial suspension as previously described (Yin *et al*., 2020). Briefly, conidia of *F. graminearum* PH-1 were harvested from cultures on 5-day-old carboxymethyl cellulose medium and adjusted to 10^5^ spores/ml in sterile water with/without 6 μM AXD. The susceptible wheat variety Xiaoyan 22 was inoculated by injecting 10 µl of spore suspension into the florets at the central section of the wheat spikelet. After inoculation, the wheat heads were covered with plastic bags to maintain humidity. The bags were removed at 3 dpi. Disease symptoms were recorded at 14 dpi.

### Accession numbers

The genes from this article can be found in the GeneBank database under the following accession numbers: MoGep4 (XP_003712260.1), MoCld1 (XP_003713872.1), MoTaz1 (XP_003713659.1), Pkc1 (XP_003719292.1), and Mps1 (XP_003712437.1).

## Results

### *MoGEP4* is involved in development, conidiation and stress response in *M. oryzae*

To investigate the function of CL in *M. oryzae*, we used homologous recombination to generate mutants for genes homologous to yeast *TAM41*, *PGS1*, *GEP4* and *CRD1* that are involved in *de novo* CL biosynthesis as well as genes homologous to yeast *CLD1* and *TAZ1* that catalyze CL remodeling (Schlame and Greenberg, 2017) (Figure S1a). We obtained null mutants for three genes, including *MoGEP4, MoCLD1* and *MoTAZ1* (Figure S1B). Compared to the wild-type (WT) strain and complementation strain (CP1), only the *Mogep4* mutant (M60) grew more slowly on complete medium (CM) and oatmeal tomato agar (OTA) plates (Figure 1a, b). On TB3 plates, which are commonly usually used for protoplast regeneration in *M. oryzae* with special osmotic homeostasis, the growth rate of *Mogep4* was partially rescued to a level 75% of the WT (Figure 1a). We examined conidiation in the WT, *Mogep4*, *Mocld1*, *Motaz1*, and CP1 strains, and observed that *Mogep4* was significantly reduced in conidiation with abnormal conidial morphology. In contrast, conidiation of the *Mocld1* and *Motaz1* mutants was comparable to the WT (Figure 1c, d).

**Figure 1.**
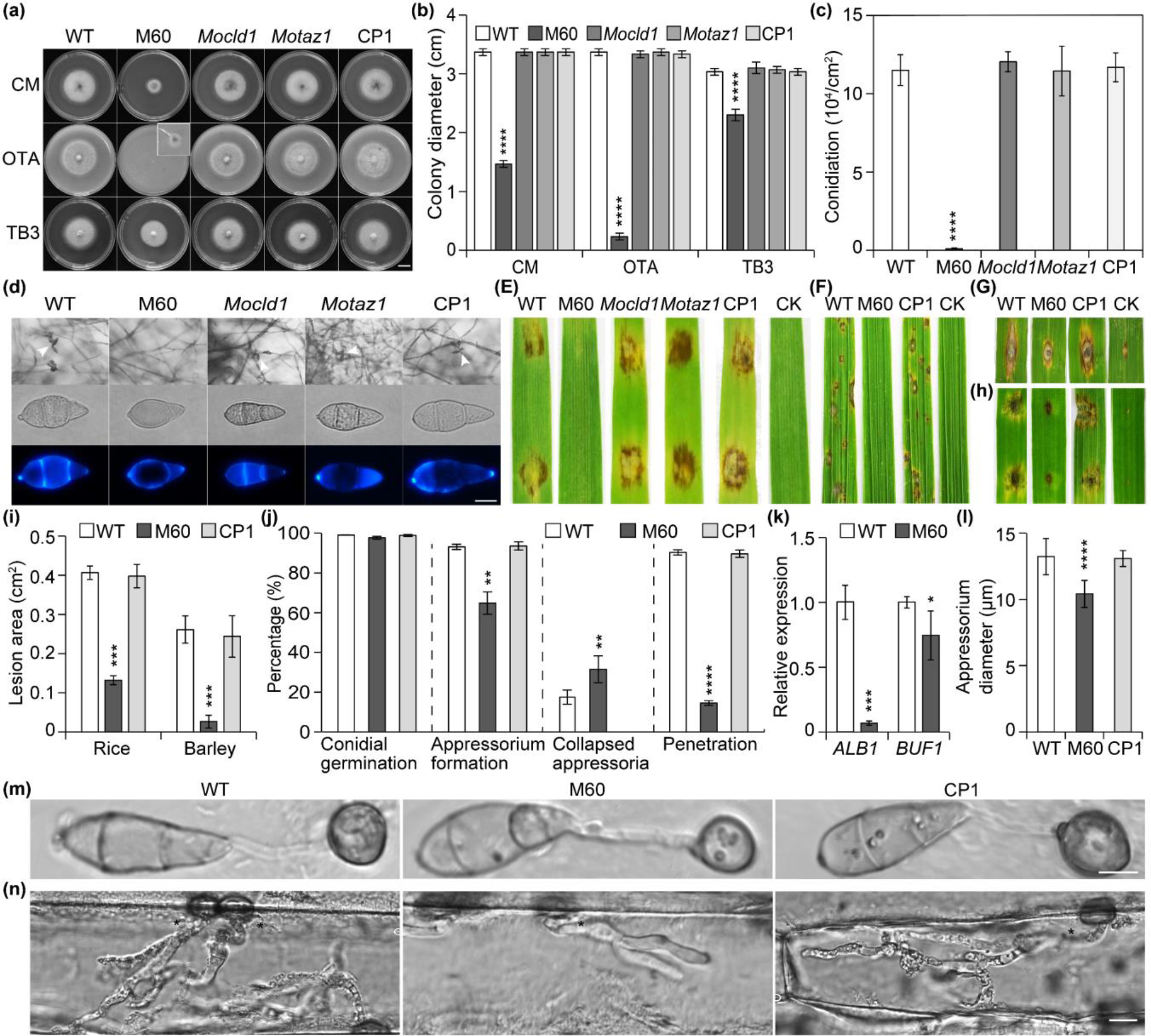
*MoGEP4* is required for growth, conidiation, pathogenicity, and infection-related morphogenesis in *M. oryzae*. **(a)** Five-day-old cultures of the wild-type (WT), *Mogep4* mutant (M60), *Mocld1* mutant, *Motaz1* mutant, and *Mogep4* complementation (CP1) strains on complete medium (CM), oatmeal tomato agar (OTA), and TB3 plates. The inset of the M60 strain on the OTA plate is a close-up view of the colony, Bar, 1 cm. **(b)** Colony diameters of strains shown in **(a)** on CM, OTA, and TB3 plates. **(c)** Conidiation of the WT, M60, *Mocld1* mutant, *Motaz1* mutant, and CP1 strains. **(d)** The *Mogep4* mutant is defective in sporulation and conidial morphology. Calcofluor white (CFW) was used to stain the conidial cell wall and septa. Arrows indicate conidia in the top panel. Bar, 10 µm. **(e)** Drop-inoculation of the WT, M60, *Mocld1*, *Motaz1*, and CP1 strains on barley leaves at 5 days post-inoculation (dpi). Water was used as the control (CK). The susceptible intact rice seedlings **(f)**, punched rice **(g)** and barley leaves **(h)** were inoculated with conidial suspensions of the WT, M60, and CP1 strains and photographed at 7 dpi. **(i)** Analysis of lesion area on rice and barley samples shown in **(g, h)**. **(j)** Percentages of conidial germination, appressorium formation, collapsed appressorium (appressorium turgor was measured with the incipient cytorrhysis assay in 2 M glycerol), and penetration. **(k)** qRT-PCR assays of melanin biosynthesis genes *ALB1* and *BUF1*. **(l)** Appressorium diameters of the WT, M60, and CP1 strains on onion epidermis at 30 hpi. At least 30 appressoria were analyzed for each strain. **(m)** Appressorial morphology on onion epidermis at 30 hpi. Bar, 10 µm. **(n)** Invasive hyphae of the WT, M60, and CP1 strains on barley leaves at 30 hpi. Asterisks indicate appressoria. Bar, 10 µm. Error bars in **(b, c, i, j, k, l)** indicate standard deviations, and asterisks indicate statistically significant differences using the unpaired Student’s *t*-test (\**P* < 0.05, \*\**P* < 0.01, \*\*\**P* < 0.001, \*\*\*\**P* < 0.0001).

We further examined aerial hyphal growth of *Mogep4* and observed that aerial hyphal growth of *Mogep4* was significantly decreased (Figure S2a), and the apical cells of hyphae were shorter compared to the WT and CP1 strains (Figure S2b, c). In addition, we observed that *Mogep4* was hypersensitive to oxidative stress.

However, under salt and osmotic stress, *Mogep4* grew faster than the WT and CP1 strains (Figure S2d, e), indicating that CL may play a role in osmotic stress balance. These results collectively demonstrate that *MoGEP4* regulates vegetative growth, aerial hyphal growth, conidiation, and stress responses in *M. oryzae*.

### *MoGEP4* is required for pathogenicity and infection-related morphogenesis in *M. oryzae*

In plant infection assays, mutant *Mogep4* caused no lesions on detached barley leaves at five days post-inoculation (dpi), whereas abundant typical necrotic lesions developed on leaves inoculated with WT, *Mocld1*, *Motaz1*, and *Mogep4* complementation strains (Figure 1e). Because only mutant *Mogep4* is involved in pathogenicity, we focused on the *MoGEP4* gene in subsequent studies. Similarly, in spray inoculation assays on seedlings of the susceptible rice cultivar LTH (Yang *et al*., 2022), *Mogep4* failed to develop lesions at 6 dpi, whereas the control strains caused abundant typical necrotic blast lesions (Figure 1f). On wounded rice and barley leaves, the lesion size caused by *Mogep4* was much smaller compared to that caused by the WT strain (Figure 1g-i). We further measured the rates of conidial germination, appressorium formation, and turgor pressure in *Mogep4*. While there was no difference in conidial germination between *Mogep4* and the WT strain (Figure 1j), the appressorium formation rate in *Mogep4* was reduced to 65% versus 90% in the WT strain at 24 hpi (Figure 1j). Similar results were observed for the turgor pressure within appressoria of *Mogep4*; more appressoria were collapsed in *Mogep4* than the WT under osmotic stress of 2 M glycerol (Figure 1j). Furthermore, expression levels of melanin biosynthesis-related genes *ALB1* and *BUF1* were significantly lower in the *Mogep4* mutant than in the WT strain (Figure 1k), consistent with the weakly melanized appressoria in *Mogep4* (Figure 1m). Additionally, the diameter of appressoria formed by *Mogep4* was smaller than that of the WT and CP1 strains (Figure 1l, m). We also observed that both lipid and glycogen metabolism were severely delayed in *Mogep4*. The lipid droplets and glycogen were still observed in the appressorium of the *Mogep4* mutant but not detected in the WT strain at 24 hours post inoculation (hpi) (Figure S3). In penetration assays, we found that only 15% of *Mogep4* appressoria penetrated the host cells at 30 hpi compared to the 90% penetration rate of the WT strain (Figure 1j, n). The invasive growth rate of *Mogep4* was much slower than the WT and CP1 strains. These results indicate that MoGep4 is required for penetration and invasive growth in *M. oryzae*.

### MoGep4 functions in CL biosynthesis

MoGep4, homologous to *Saccharomyces cerevisiae* Gep4 with a 44% protein identity, is predicted to be a conserved phosphatase (Figure 2a). Therefore, we used the yeast *gep4* mutant to study the possible biochemical function of MoGep4. When expressed in yeast, *MoGEP4* complemented the *gep4* mutant, rescuing the growth of *gep4* at 37°C (Figure 2b). To further analyze the function of MoGep4 in *M. oryzae*, we isolated mitochondria from protoplasts and analyzed phospholipids, including CL, by two-dimensional thin layer chromatography (2D-TLC). The CL level was significantly reduced in the *Mogep4* mutant, which was confirmed by the LC-MS/MS lipidomics results (Figure 2c, d). These results suggested that the MoGep4 protein plays a role similar to that in yeast in CL biosynthesis.

**Figure 2.**
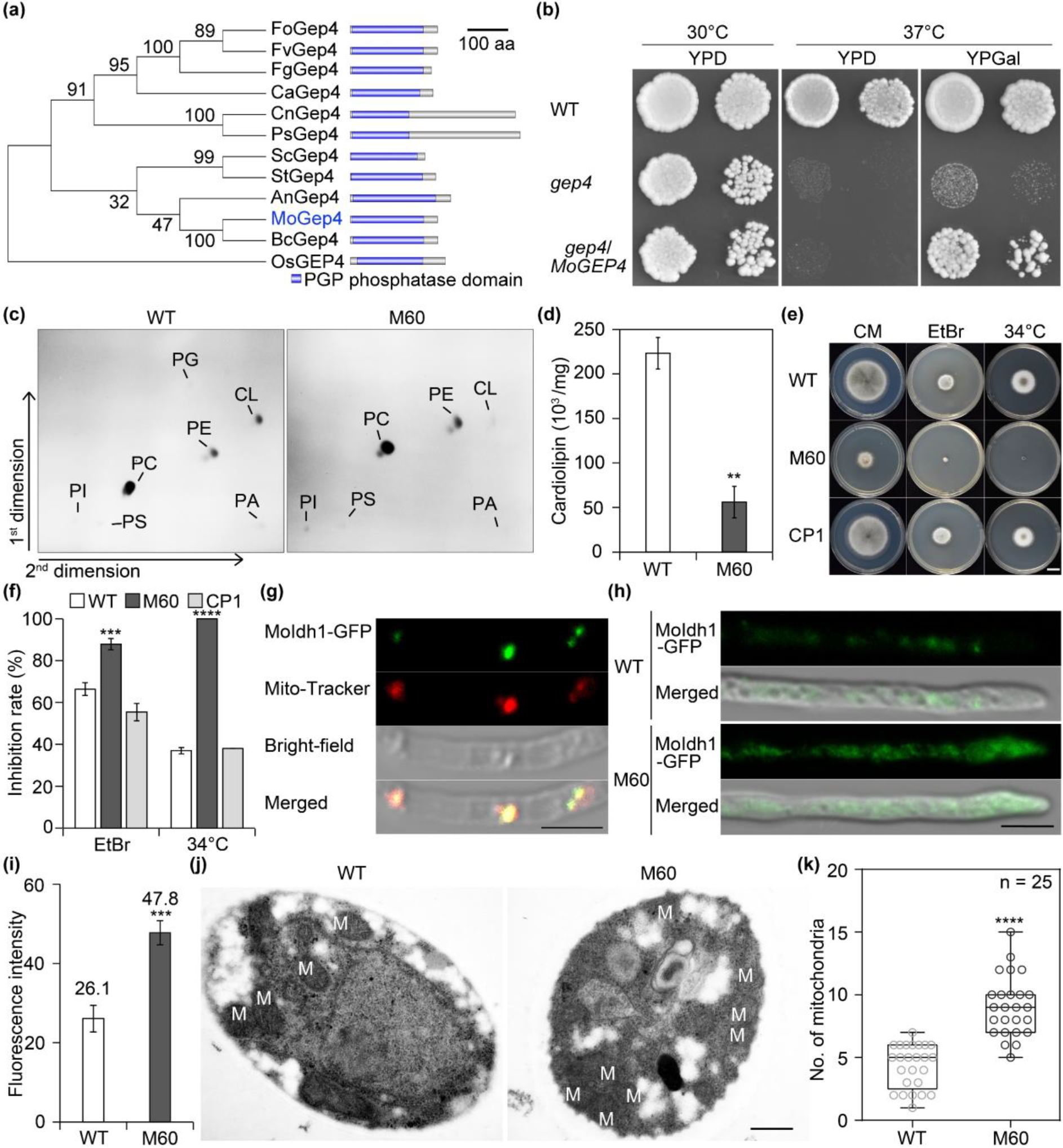
*MoGEP4* is involved in cardiolipin (CL) biosynthesis and mitochondrial function in *M. oryzae*. **(a)** Phylogenetic analysis of MoGep4 and its orthologs by the neighbor-joining method with MEGA 7 (left panel), including *Aspergillus nidulans* (AnGep4), *Botrytis cinerea* (BcGep4), *Candida albicans* (CaGep4), *Cryptococcus neoformans* (CnGep4), *Fusarium graminearum* (FgGep4), *Fusarium oxysporum* (FoGep4), *Fusarium verticillioides* (FvGep4), *Oryza sativa* (OsGEP4), *Puccinia striiformis* (PsGep4), *Saccharomyces cerevisiae* (ScGep4), and *Septoria tritici* (StGep4). Values at the branching points represent the results of bootstrap analysis. Schematic representation of the domains of Gep4 proteins (right panel). **(b)** Expression of *MoGEP4* rescued the growth defect of the yeast *gep4* mutant. Yeast strains BY4741 (WT), the *gep4* mutant (*gep4*), and transformants of the *gep4* mutant carrying pYES2-*MoGEP4* (*gep4*/*MoGEP4*) were cultured on YPD and YPGal plates at 30°C and 37°C for 4 days, respectively. **(c)** CL content is reduced in the *Mogep4* mutant. Mitochondrial phospholipids of the WT and *Mogep4* mutant (M60) strains were analyzed by two- dimensional thin-layer chromatography (2D-TLC). PA, phosphatidic acid; PC, phosphatidylcholine; PE, phosphatidylethanolamine; PG, phosphatidylglycerol; PI, phosphatidylinositol; PS, phosphatidylserine. **(d)** Lipidomics analysis of CL in the WT and M60 strains. Six-day-old cultures **(e)** and growth inhibition rates **(f)** of the WT, M60, and CP1 strains on CM plates supplemented with/without 2.5 µg/ml Ethidium Bromide (EtBr) or at 34°C. **(g)** The fluorescent signal of MoIdh1-GFP is colocalized with Mito-Tracker in hyphae. Bars, 5 µm Distribution **(h)** and average fluorescent signal intensity **(i)** of mitochondrion-localized MoIdh1- GFP in the WT and M60 strains (more than 30 hyphae were examined). Bar, 5 µm. The GFP signal intensity was analyzed with ImageJ. **(j)** Mitochondrial morphology of the WT and M60 strains under transmission electron microscopy. The strains were cultured in the MM-N for 2 h for induction of mitophagy before sampling. Bar, 0.5 µm. **(k)** The number (No.) of mitochondria was determined in the WT and M60 strains. Error bars in **(d, f, i, k)** indicate standard deviations, and asterisks indicate statistically significant differences using the unpaired Student’s *t*-test (\*\**P* < 0.01, \*\*\**P* < 0.001, \*\*\*\**P* < 0.0001).

We expressed MoGep4-GFP fusion protein in *Mogep4* as the CP1 strain. The fusion protein rescued the defects of *Mogep4* (Figure 1 and Figure S2), indicating that the MoGep4-GFP fusion protein is fully functional. We observed fluorescent signals of MoGep4-GFP in hyphae (Figure S4) and found that it co- primarily localized with the mitochondrial dye Mito-Tracker. Therefore, our results together demonstrate that MoGep4 functions as a mitochondrial protein that is required for CL biosynthesis in *M. oryzae*.

### Loss of *MoGEP4* leads to defects in mitochondrial function

To determine whether loss of *MoGEP4* affects mitochondrial functions, we inoculated the WT and *Mogep4* mutant on CM plates at 34°C and at 28°C on CM supplemented with ethidium bromide (EtBr), which is known to induce the loss of mitochondrial DNA (Osman *et al*., 2010). Interestingly, we found that *Mogep4* was incapable of growth at 34°C (Figure 2e, f). In the presence of EtBr (2.5 ug/ml), the growth rate of *Mogep4* was substantially reduced compared to the WT strain at 28°C (Figure 2e, f), suggesting that MoGep4 is involved in mitochondrial stress responses in *M. oryzae*.

We used the mitochondrion-specific protein MoIdh1 (Figure 2g), an ortholog of yeast Idh1 that has been used as a mitophagy marker (Yamashita *et al*., 2016), to analyze mitochondrial distribution and abundance in *Mogep4*. Microscopic results showed that accumulation in mitochondria of *Mogep4* was higher than in the WT strain (Figure 2h, i). Similar results were observed under transmission electron microscopy (Figure 2j, k). Given that the growth of the *Mogep4* mutant is significantly inhibited on OTA plates that are nutrient poor, and that mitochondria are important for energy production, these results together suggest that MoGep4 is critical for mitochondrial function, and that the ability of clearing excessive mitochondria may be impaired in *Mogep4*.

### *MoGEP4* is essential for mitophagy in *M. oryzae* infection

Because more mitochondria were observed in the *Mogep4* mutant than in the WT strain, we hypothesized that loss of CL leads to defects in mitophagy. We investigated the mitophagy process during nitrogen starvation. The GFP fluorescence signal of MoIdh1 in the WT strain was observed in the vacuole after mitophagy induction for 0.5 h, but was not detected in *Mogep4* (Figure 3a). At 12 h, GFP fluorescence signals were observed in the vacuole of the *Mogep4* mutant as well as the WT, which suggests that mitophagy is delayed in *Mogep4*, consistent with the result that the MoIdh1-GFP degradation process was notably reduced in *Mogep4* and confirming that *Mogep4* is impaired in mitophagy (Figure 3b). We further found that the MoIdh1-GFP signal was observed in the vacuoles and invasive hyphae of the WT strain at 48 hpi but were not detected in the vacuoles of *Mogep4* (Figure 3c), suggesting that mitophagy is impaired during the infection process of *Mogep4*.

**Figure 3.**
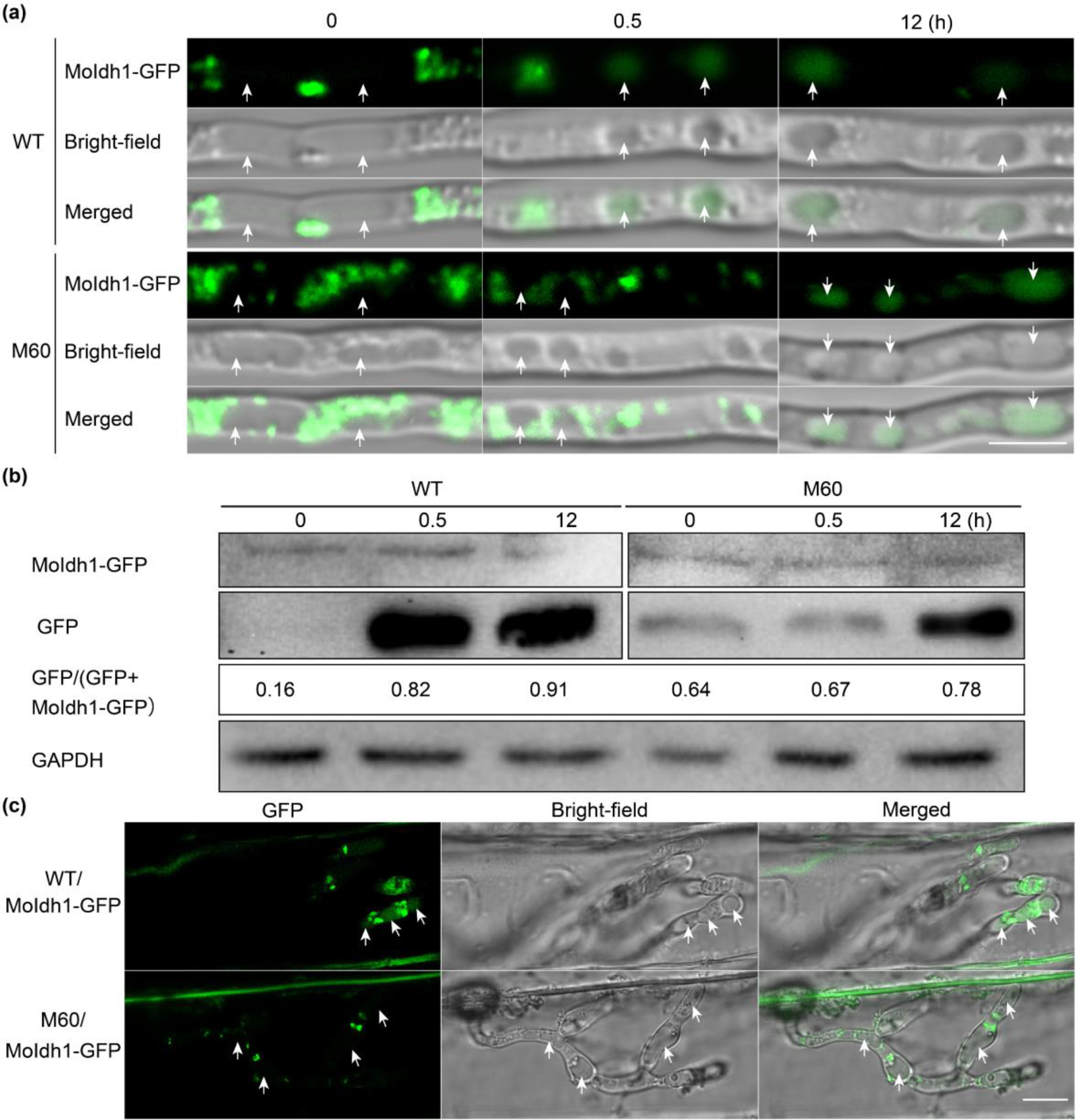
*MoGEP4* is essential for mitophagy-mediated infection. **(a)** MoGep4 is required for mitophagy in invasive hyphae. The WT and *Mogep4* (M60) strains expressing *MoIDH1*-GFP were inoculated onto barley leaves and photographed at 48 hpi. Arrows indicate vacuoles. Bar, 10 µm. **(b)** The *Mogep4* mutant is impaired in mitophagy. The same set of strains shown in **(a)** was induced for mitophagy. Arrows indicate vacuoles. Bar, 10 µm. **(c)** The *Mogep4* mutant is reduced in mitophagy. Immunoblotting analysis of mitophagy in the WT and M60 strains, indicated by the ratio of free GFP to total GFP. Glyceraldehyde 3-phosphate dehydrogenase (GAPDH) indicates the protein loading for each lane.

### MoGep4 is involved in Mps1-MAPK activation

To investigate the role of MoGep4 in the Mps1-MAPK signaling pathway, we examined the growth rate of *Mogep4* under different cell wall stressors. Mutant *Mogep4* was more sensitive to cell wall stressors Congo Red and Calcofluor White than the WT and CP1 strains (Figure 4a, b). Our results also showed that *Mogep4* was more sensitive to cell-wall-digesting enzyme than the WT (Figure 4c). In accord, Mps1 phosphorylation was significantly reduced in *Mogep4* compared to the WT and CP1 strains under the mitophagy conditions (Figure 4d). Interestingly, the phosphorylation level of Mps1 was significantly increased in the *MoGEP4* overexpression strain GOX1, suggesting that MoGep4 plays a regulatory role in the activation of the Mps1- MAPK pathway. Ceramides have been reported to activate Mps1-MAPK (Huwiler *et al*., 1996, Fox *et al*., 2007, Liu *et al*., 2019). Therefore, we hypothesized that MoGep4 affects the level of ceramides to regulate the Mps1-MAPK signaling pathway (Figure 4e). Indeed, the sphingolipidomics assays showed that ceramide levels were remarkably reduced in *Mogep4* compared to the WT strain (Figure 4f). These data support a conclusion that MoGep4 plays a critical role in the biosynthesis of ceramides and activation of the Mps1- MAPK pathway in *M. oryzae*.

**Figure 4.**
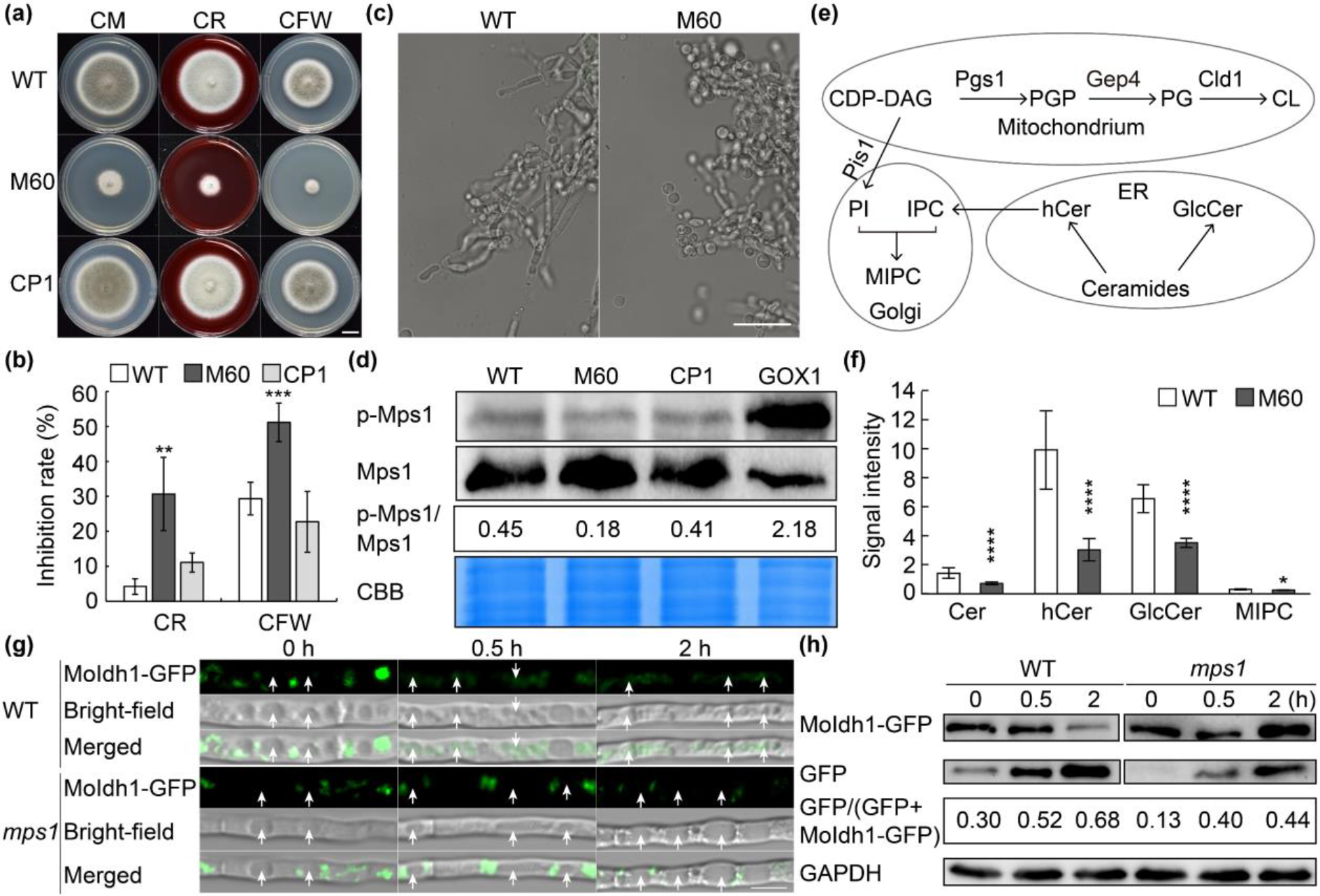
The *Mogep4* mutant is defective in the cell wall integrity signaling pathway. Six-day-old cultures **(a)** and inhibition rates **(b)** of the wild-type (WT), *Mogep4* mutant (M60), and complementation (CP1) strains on CM plates supplemented with different stressors, including Congo Red (CR, 200 µg/ml) and Calcofluor White (CFW, 200 µg/ml). Bar, 1 cm. **(c)** The *Mogep4* mutant is hypersensitive to cell-wall-digesting enzymes. Mycelia of the strains were photographed 30 min after enzyme treatment. Bar, 25 µm. **(d)** Mps1 activation is reduced in the *Mogep4* mutant. Immunoblotting of the WT, M60, CP1, and *MoGEP4* overexpression (GOX1) strains using Mps1 and pospho-p44/42 antibodies. The proteins were isolated from mycelia cultured in MM-N for 2 h. **(e)** The simplified pathway of phospholipid biosynthesis involved in sphingolipids in yeast. Cer, free ceramides with non-hydroxy fatty acids; hCer, free ceramides with 2-hydroxy fatty acids; GlcCer, monoglucosyl ceramides; MIPC, mannosyl-inositol-phospho- ceramides. **(f)** Ceramides are reduced in the *Mogep4* mutant. Normalized signal intensity represents the sphingolipid levels. **(g)** The *mps1* mutant is defective in mitophagy. The hyphae of the WT and *mps1* expressing the *MoIDH1*-GFP were cultured in MM-N and photographed at 0.5 and 2 hpi, respectively. **(h)** The *mps1* mutant is reduced in mitophagy. Immunoblotting analysis of mitophagy in the WT and *mps1* strains, indicated by the ratio of free GFP to total GFP. GAPDH denotes the protein loading for each lane. Arrows indicate vacuoles. Bar, 10 µm. Error bars in **(b and f)** indicate standard deviations, and asterisks indicate statistically significant differences using the unpaired Student’s *t*-test (\**P* < 0.05, \*\**P* < 0.01, \*\*\**P* < 0.001, \*\*\*\**P* < 0.0001).

Because both mitophagy and Mps1 phosphorylation are altered in the *Mogep4* mutant, Mps1 and mitophagy may also be associated. We tested this hypothesis in the *mps1* mutant and observed that MoIdh1- GFP signals were observed in the vacuoles of the WT strain but not in the *mps1* mutant (Figure 4g, h), showing that Mps1 is involved in mitophagy.

### AXD inhibits the enzyme activity of MoGep4

The homolog of MoGep4, PTPMT1, is targeted by alexidine dihydrochloride (AXD) (Doughty-Shenton *et al*., 2010). Here, for future fungicide development, we tested whether AXD can be used to inhibit MoGep4 enzyme activity. First, we computationally analyzed whether AXD could bind to MoGep4. Molecular docking using the MoGep4 structure predicted by AlphaFold2 was performed, and the results showed that Asp60 of MoGep4 is a potential residue that interacts with AXD via hydrogen bonds, with a binding energy at -5.6 kcal/mol (Figure 5a). We then performed Surface Plasmon Resonance (SPR) assays and found that AXD binds to MoGep4 with a dissociation equilibrium constant (*KD*) of 4.26 µM (Figure 5b, c). Further thermal shift assays showed that the thermal transition temperature (Tm) of MoGep4 changed 0.8°C compared with the control, higher than 0.4°C, indicating that AXD attenuated the structural stability of MoGep4 (Figure 5d). These results together showed that AXD binds to MoGep4.

**Figure 5.**
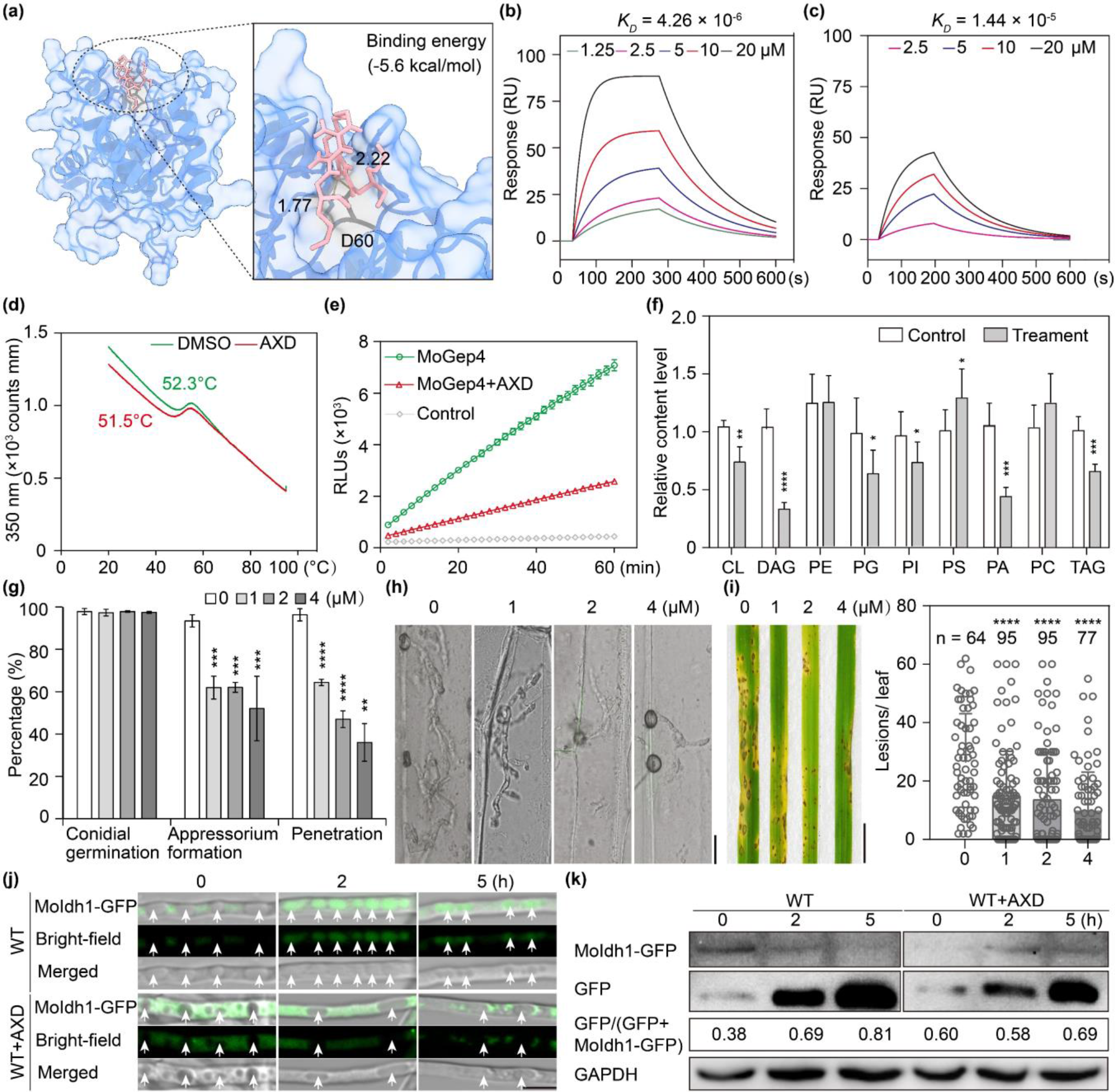
Alexidine dihydrochloride (AXD), a MoGep4-interacting chemical, suppresses fungal infection. **(a)** Molecular docking of MoGep4 (blue) with AXD (pink). The close-up view shows the predicted hydrogen bonds, represented by dashed lines. SPR analysis of the binding affinity of AXD to MoGep4-GST **(b)** and the GST tag as the control **(c)**. **(d)** Differential scanning fluorimetry using intrinsic protein fluorescence (nano DSF) scans for MoGep4 treated with 5% DMSO (red) or 50 µm AXD (green) at 350 nm. **(e)** AXD inhibits MoGep4 enzyme activity. The GST tag was used as the control. **(f)** Lipidomics assays of *M.oryzae* treated with AXD (4 μM). **(g)** Conidial germination, appressorium formation, and penetration assays of *M. oryzae* supplemented with/without AXD. **(h)** Invasive growth of *M. oryzae* in barley was inhibited by AXD. Photos were taken at 30 hpi. Bar, 10 µm. **(i)** Infection assays with conidial suspensions supplemented with/without AXD. Three-week-old rice seedlings were used in the assay. Scale bar, 1 cm. **(j)** AXD inhibits mitophagy in *M. oryzae*. MoIdh1-GFP was used as the mitophagy marker. Arrows indicate vacuoles. Bar, 10 µm. **(k)** Immunoblotting analysis of mitophagy in *M. oryzae* treated with AXD, indicated by the ratio of free GFP to total GFP. GAPDH denotes the protein loading for each lane. Means and standard deviations were calculated from three replicates, and asterisks indicate statistically significant differences using the Student’s *t*-test (\**P* < 0.05, \*\**P* < 0.01, \*\*\**P* < 0.001, \*\*\*\**P* < 0.0001).

To investigate whether AXD inhibits the enzyme activity of MoGep4, MoGep4 was expressed in *Escherichia coli* and purified. AXD was added to the enzyme reaction buffer, and the results showed that the enzyme activity of MoGep4 was significantly inhibited by 62.9% by AXD at 4 µM (Figure 5e). We subsequently tested whether AXD inhibits CL biosynthesis *in vivo* by adding AXD to the fungal culture. The lipidomics results showed that AXD significantly decreased PG and CL levels by 39.5% and 29.4%, respectively, in *M. oryzae* (Figure 5f), indicating that AXD suppresses MoGep4 enzyme activity and inhibits CL biosynthesis in *M. oryzae*.

To analyze the function of the predicted AXD-binding residue of MoGep4, we mutated the residue and transformed the mutant allele *MoGEP4^D60A^* into the yeast *gep4* mutant. When expressed in yeast, *MoGEP4* but not the *MoGEP4^D60A^* mutant allele complemented the *gep4* mutant, by rescuing the growth of *gep4* at 37°C (Figure S5a). Similarly, in *M. oryzae*, *MoGEP4^D60A^* could not rescue the growth and pathogenicity of the *Mogep4* mutant (Figure S5b, c, e). Furthermore, the *Mogep4^D60A^* mutant (*MoGEP4^D60A^*) was reduced in its sensitivity to AXD, similar to the *Mogep4* mutant (Figure S5b, c), and SPR results showed that the binding affinity is also reduced (Figure 5b, c and S5d). These results indicated that Asp60 is important for AXD binding and MoGep4 enzyme activity.

### AXD is effective in the control of crop diseases in the field

To verify whether AXD can inhibit infection-related processes, we treated *M. oryza* with 4 μM AXD, which efficiently inhibited appressorium formation, appressorium penetration, invasive hyphal growth and plant infection but not conidial germination of *M. oryzae* (Figure 5g-i). Our results further showed that AXD significantly inhibited mitophagy in *M. oryzae* (Figure 5j, k).

Because Gep4 proteins are conserved in filamentous plant pathogens, and similar protein structures and binding pockets were observed in Gep4 homologs of different pathogens (Figure S6), we further tested AXD on nine other pathogens including *Botrytis cinerea*, *Bipolaris maydis*, *Bipolaris sorokiniana, Fusarium graminearum*, *Leptosphaeria biglobosa*, *Monilinia fructicola, Phytophthora sojae*, *Valsa mali*, and *Valsa pyri*. The inhibition rates on these nine pathogens by AXD at 4 μM ranged from 9% to 79%, with *L. biglobosa* the most sensitive (Figure 6a, b). We next tested AXD on plant infection by *B. cinerea*, *V. pyri* and *M. oryzae*, and the results showed that AXD efficiently inhibited plant infection by these pathogens (Figure 6c-f), without visible side effects on the plants (Figure S7). Finally, we carried out field trials to evaluate the efficacy of AXD on rice blast and Fusarium head blight of wheat and observed that AXD reduced the severity of rice blast at a concentration of 6 μM (Figure 6g, h). For FHB, the infection was completely inhibited when the inoculum was supplemented with AXD at 6 μM (Figure 6i, j). Taken together, our results showed that AXD displays broad-spectrum antifungal activity under both the greenhouse and field conditions.

**Figure 6.**
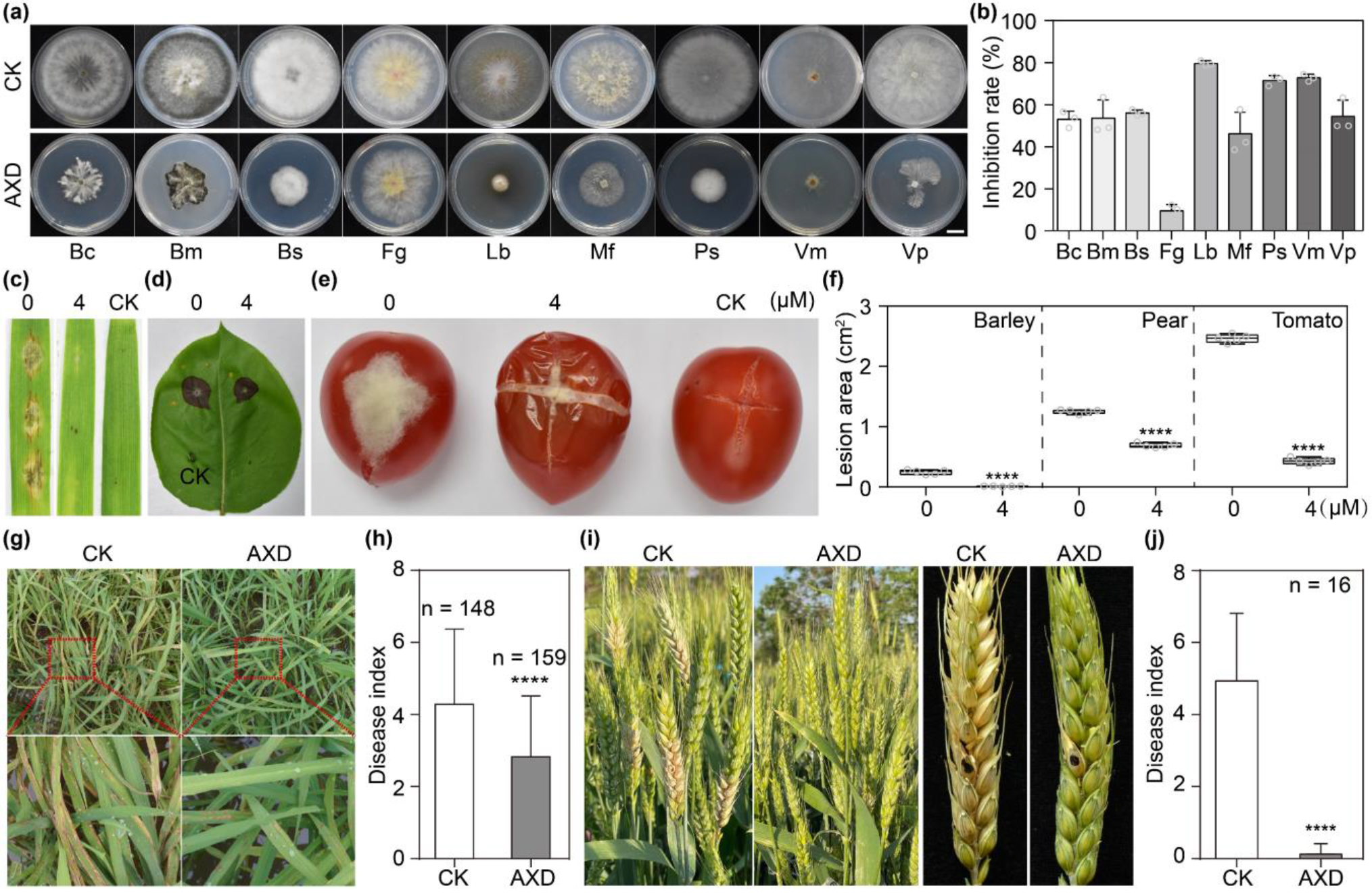
Alexidine dihydrochloride (AXD) shows broad-spectrum antifungal activity. Inhibitory effects **(a)** and inhibition rates **(b)** of AXD (4 μM) on different pathogens grown on plates supplemented with/without AXD. Bc, *B. cinerea*; Bm, *B. maydis*; Bs, *B. sorokiniana*; Fg, *F. graminearum*; Lb, *L. biglobosa*; Mf, *M. fructicola*; Ps, *P. sojae*; Vm, *V. mali*; and Vp, *V. pyri*. Bar, 1 cm. **(c-e)** Plant infection assays of different pathogens with/without 4 μM AXD. **(f)** Lesion areas in barley, pear, and tomato samples shown in **(c-e)**. The lesion area was measured at 5 dpi. **(g)** Field trials of AXD on rice. AXD (6 µM) was sprayed onto rice seedlings at 30 days post-planting. The solvent dimethyl sulfoxide (DMSO) was used as the control. **(h)** Rice blast symptoms were examined at 16 dpi. **(i)** Effect of AXD on Fusarium head blight in the field. **(J)** Representative infected wheat heads and the disease index of Fusarium head blight at 14 dpi. Means and standard deviations were calculated from three replicates, and asterisks indicate statistically significant differences using the unpaired Student’s *t*-test (\*\*\*\**P* < 0.0001).

## Discussion

Fungal virulence factors are potential targets for controlling crop diseases (Liu *et al*., 2024). In this study, we sought to understand the role of enzymes involved in CL biosynthesis in fungal pathogenesis and explored the possibility of targeting CL biosynthetic enzymes for fungicide development. We demonstrated that CL biosynthesis-related gene *MoGEP4* is involved in fungal growth, conidiation, virulence, mitophagy, and Mps1-MAPK activation in *M. oryzae*, and that AXD inhibits MoGep4 activity and effectively controls crop diseases in the field.

Our results showed the conserved and diverse roles of CL biosynthetic enzymes. *MoGEP4* complemented the defects of yeast *gep4* mutant, indicating the conserved functions of MoGep4 in yeast. The *Mogep4* mutant is defective in mitochondrial functions and shows hypersensitivity to mitochondrial stressors, confirming conserved roles of CL in mitochondrial function and homeostasis (Ikon and Ryan, 2017).

However, mutation of *MoTAZ1* and *MoCLD1*, CL-remodeling genes, caused no obvious defects in growth, conidiation or virulence, which contrasts with the results in yeast, in which *TAZ1* and *CLD1* are critical for mitochondrial functions and vegetative growth (Beranek *et al*., 2009, Baile *et al*., 2014). In humans, the CL remodeling enzyme tafazzin encoded by *TAZ1*, is involved in the life-threatening genetic disorder Barth syndrome (Oemer *et al*., 2022). Taken together, *GEP4* homologs, involved in CL biosynthesis, play conserved roles in different species but not the CL-remodeling-related genes, *MoTAZ1* and *MoCLD1*.

In *M. oryzae*, *MoGEP4* is involved in mitophagy and Mps1-MAPK activation through different mechanisms. Both *Mogep4* and *Moatg24*, a mitophagy-defective mutant in *M. oryzae* (He et al., 2013), are impaired for fungal growth, conidiation and virulence, indicating the association between MoGep4 and mitophagy. These resulting phenotypes are likely because *MoGEP4* is required for CL biosynthesis in the mitochondrial membrane, which is important for normal mitochondrial structures, mitophagy and fungal infection (Shen *et al*., 2017, Kou *et al*., 2019, Xiao *et al*., 2021). Both the *Mogep4* mutant and mutants in the Mps1-MAPK pathway are hypersensitive to cell wall stressors, showing reduced conidiation and osmotically remedial phenotypes (Jagernath *et al*., 2021) (Xu *et al*., 1998, Heinisch *et al*., 1999, Flández *et al*., 2004, Yin *et al*., 2016). The involvement of MoGep4 in the Mps1-MAPK pathway is likely through ceramides (Huwiler *et al*., 1996, Fox *et al*., 2007), which induce Mps1 phosphorylation and are required for pathogenesis of *M. oryzae* and *F. graminearum* (Rittenour *et al*., 2011, Liu *et al*., 2019) but are reduced in the *Mogep4* mutant.

Taken together, *MoGEP4* is involved in the regulation of mitophagy, the Mps1-MAPK signaling pathway and pathogenesis.

Our results showed that AXD efficiently inhibited fungal infection and controlled rice blast and FHB in the field. AXD targets MoGep4 in *M. oryzae*, supported by multiple lines of evidence including molecular docking, SPR, nanoDSF and enzyme activity assays. Secondly, most phenotypes caused by AXD are reminiscent of those of the *Mogep4* mutant, including reduced growth and conidiation, appressorium formation, attenuated virulence, and altered phospholipid levels. However, the *M. oryzae* phenotypes resulting from AXD treatment are broader than those of the *Mogep4* mutant. For example, AXD-treated mycelia of *M. oryzae* are less pigmented compared to the *Mogep4* mutant, suggesting that AXD could have multiple targets. Fungicides with multiple targets are not uncommon. In fact, many fungicides have a multi-target mode of action, which enhances the efficacy of the fungicide and reduces the development of fungicide resistance. For example, chrysomycin A has a multiple target mode of action, which competitively inhibits GlmU and DapD, targeting the biosynthetic pathways of peptidoglycan and lysine precursor in *Staphylococcus aureus*, respectively (Jia *et al*., 2023). Further research is needed to investigate the antifungal mechanism of AXD, which provides information for rational modification of AXD to enhance its fungicidal activity.

Multiple strategies can be used to reduce the possible side effects of AXD in disease control. Chemical modification is an effective method for generating highly specific and low-toxicity AXD devivatives. For example, Shen et al. (2017) improved the therapeutic efficacy of phosphorothioate antisense oligonucleotides and reduced toxicity through chemical modifications (Hu *et al*., 2023). Alternatively, using nanomaterials to deliver fungicides is an effective strategy to reduce the possible side effects (Balusamy *et al*., 2023). For example, Fischer et al. (2019) used nanocarriers to deliver Pyraclostrobin to plants for controlling grapevine fungal disease and minimize the side effects by releasing only the fungicide encased within the nanocarrier upon degradation by fungal enzymes. Finally, the working concentration of AXD in disease control is only about one-tenth of that showing toxicity to mammalian cell lines (Doughty-Shenton *et al*., 2010), not to mention that the minimal amount of AXD can reach the worker. Last but not least, these strategies can be integrated to enhance the effective and environment-friendly use of AXD in agriculture.

In summary, this study demonstrated that MoGep4 is critical for CL biosynthesis, mitophagy, MAPK activation, and virulence in *M. oryzae*. Our results also showed that AXD inhibits MoGep4 enzyme activity, displays broad-spectrum antifungal activity against 10 pathogens and is effective in disease control in the field, a promising fungicide candidate against a wide range of crop diseases.

## Acknowledgements

We thank Drs. Larry D. Dunkle and Charles P. Woloshuk (Purdue University), Dr. Lianhu Zhang (Jiangxi Agriculture University) for critical reading of this manuscript. We thank Dr. Cong Jiang (Northwest A&F University) and Dr. Junfeng Liu (China Agricultural University) for fruitful discussions. At Huazhong Agricultural University, we thank Prof. Chunyu Zhang for technical assistance in 2D-TLC analysis and we thank Dr. X. Shen for the support of data acquisition and analysis. We thank Ms. Dongqin Li from the National Key Laboratory of Crop Genetic Improvement for assistance in mass spectrometry analysis. Multiple microscopic data were acquired at the National Key Laboratory of Agricultural Microbiology Core Facility.

This work was also supported by Hubei Hongshan Laboratory. This work was supported by the National Natural Science Foundation of China (32172373, 31801723), National Key R&D Program of China (2022YFA1304402) and Fundamental Research Funds for the Central Universities (2023ZKPY002, 2662023PY006, AML2023A05) to G. L. This work was supported by the China Postdoctoral Science Foundation (2023M741298) to P. S.

## Data availability statement

The data that support the findings of this study are available from the corresponding authors upon reasonable request.

## Author contributions

G.L. and P.S. conceived and designed the experiments. P.S., Z.J., G.S., Y.Z., M.Z., X.K., Q.S., Y.L., K.L., R.B., W.H., Q.L., T.I., J.G., and G.C. performed experimental work. Z.Q. and L.Y. performed bioinformatic analysis. R. L. analyzed the molecular docking and tested some fungal strains. G.W. performed the field experiments on wheat. G.L. and P.S. wrote the manuscript with inputs of all coauthors.

## References

1. Baile, M.G., Sathappa, M., Lu, Y.W., Pryce, E., Whited, K., McCaffery, J.M., Han, X., Alder, N.N. and Claypool, S.M. (2014) Unremodeled and remodeled cardiolipin are functionally indistinguishable in yeast. J. Biol. Chem., 289, 1768–1778.

2. Balusamy, S.R., Joshi, A.S., Perumalsamy, H., Mijakovic, I. and Singh, P. (2023) Advancing sustainable agriculture: a critical review of smart and eco-friendly nanomaterial applications. J Nanobiotechnology, 21, 372.

3. Beranek, A., Rechberger, G., Knauer, H., Wolinski, H., Kohlwein, S.D. and Leber, R. (2009) Identification of a cardiolipin-specific phospholipase encoded by the gene *CLD1* (*YGR110W*) in yeast. J. Biol. Chem., 284, 11572–11578.

4. Bligh, E.G. and Dyer, W.J. (1959) A rapid method of total lipid extraction and purification. Can J Biochem Physiol, 37, 911–917.

5. Catlett, N.L., Lee, B.-N., Yoder, O.C. and Turgeon, B.G. (2003) Split-Marker recombination for efficient targeted deletion of fungal genes. Fungal Genet Rep, 50, 9–11.

6. Chen, J., Zheng, W., Zheng, S., Zhang, D., Sang, W., Chen, X., Li, G., Lu, G. and Wang, Z. (2008) Rac1 is required for pathogenicity and Chm1-dependent conidiogenesis in rice fungal pathogen *Magnaporthe grisea*. PLoS Pathog., 4, e1000202.

7. Chu, C.T., Ji, J., Dagda, R.K., Jiang, J.F., Tyurina, Y.Y., Kapralov, A.A., Tyurin, V.A., Yanamala, N., Shrivastava, I.H., Mohammadyani, D., Wang, K.Z.Q., Zhu, J., Klein-Seetharaman, J., Balasubramanian, K., Amoscato, A.A., Borisenko, G., Huang, Z., Gusdon, A.M., Cheikhi, A., Steer, E.K., Wang, R., Baty, C., Watkins, S., Bahar, I., Bayir, H. and Kagan, V.E. (2013) Cardiolipin externalization to the outer mitochondrial membrane acts as an elimination signal for mitophagy in neuronal cells. Nat cell biol, 15, 1197–1205.

8. David, A., Islam, S., Tankhilevich, E. and Sternberg, M.J.E. (2022) The AlphaFold database of protein structures: A biologist’s guide. J. Mol. Biol., 434, 167336.

9. Doughty-Shenton, D., Joseph, J.D., Zhang, J., Pagliarini, D.J., Kim, Y., Lu, D., Dixon, J.E. and Casey, P.J. (2010) Pharmacological targeting of the mitochondrial phosphatase PTPMT1. J. Pharmacol. Exp. Ther., 333, 584–592.

10. Flández, M., Cosano, I.C., Nombela, C., Martín, H. and Molina, M. (2004) Reciprocal regulation between Slt2 MAPK and isoforms of Msg5 dual-specificity protein phosphatase modulates the yeast cell integrity pathway. J. Biol. Chem., 279, 11027–11034.

11. Fox, T.E., Houck, K.L., O’Neill, S.M., Nagarajan, M., Stover, T.C., Pomianowski, P.T., Unal, O., Yun, J.K., Naides, S.J. and Kester, M. (2007) Ceramide recruits and activates protein kinase C zeta (PKC zeta) within structured membrane microdomains. J. Biol. Chem., 282, 12450–12457.

12. Gebert, N., Joshi, A.S., Kutik, S., Becker, T., McKenzie, M., Guan, X.L., Mooga, V.P., Stroud, D.A., Kulkarni, G., Wenk, M.R., Rehling, P., Meisinger, C., Ryan, M.T., Wiedemann, N., Greenberg, M.L. and Pfanner, N. (2009) Mitochondrial cardiolipin involved in outer-membrane protein biogenesis: implications for Barth syndrome. Curr. Biol., 19, 2133–2139.

13. Hamer, J.E., Howard, R.J., Chumley, F.G. and Valent, B. (1988) A mechanism for surface attachment in spores of a plant pathogenic fungus. Science., 239, 288–290.

14. He, Y., Deng, Y.Z. and Naqvi, N.I. (2013) Atg24-assisted mitophagy in the foot cells is necessary for proper asexual differentiation in *Magnaporthe oryzae*. Autophagy, 9, 1818–1827.

15. Heinisch, J.J., Lorberg, A., Schmitz, H.P. and Jacoby, J.J. (1999) The protein kinase C-mediated MAP kinase pathway involved in the maintenance of cellular integrity in *Saccharomyces cerevisiae*. Mol. Microbiol., 32, 671–680.

16. Hu, X., Yang, P., Chai, C., Liu, J., Sun, H., Wu, Y., Zhang, M., Zhang, M., Liu, X. and Yu, H. (2023) Structural and mechanistic insights into fungal β-1,3-glucan synthase FKS1. Nature, 616, 190–198.

17. Huwiler, A., Brunner, J., Hummel, R., Vervoordeldonk, M., Stabel, S., van den Bosch, H. and Pfeilschifter, J. (1996) Ceramide-binding and activation defines protein kinase c-Raf as a ceramide-activated protein kinase. Proc. Natl Acad Sci. USA., 93, 6959–6963.

18. Ikon, N. and Ryan, R.O. (2017) Cardiolipin and mitochondrial cristae organization. Biochim Biophys Acta., 1859, 1156–1163.

19. Jagernath, J.S., Meng, S., Qiu, J., Shi, H. and Kou, Y. (2021) Selective degradation of mitochondria by mitophagy in pathogenic fungi. Am. J. Mol. Biol., 11, 15–27.

20. Ji, J. and Greenberg, M.L. (2022) Cardiolipin function in the yeast *S. cerevisiae* and the lessons learned for Barth syndrome. J Inherit Metab Dis, 45, 60–71.

21. Jia, J., Zheng, M., Zhang, C., Li, B., Lu, C., Bai, Y., Tong, Q., Hang, X., Ge, Y., Zeng, L., Zhao, M., Song, F., Zhang, H., Zhang, L., Hong, K. and Bi, H. (2023) Killing of Staphylococcus aureus persisters by a multitarget natural product chrysomycin A. Sci Adv, 9, eadg5995.

22. Kou, Y., He, Y., Qiu, J., Shu, Y., Yang, F., Deng, Y. and Naqvi, N.I. (2019) Mitochondrial dynamics and mitophagy are necessary for proper invasive growth in rice blast. Mol. Plant Pathol, 20, 1147–1162.

23. Li, G., Zhang, X., Tian, H., Choi, Y.E., Tao, W.A. and Xu, J.R. (2017) MST50 is involved in multiple MAP kinase signaling pathways in *Magnaporthe oryzae*. Environ Microbiol, 19, 1959–1974.

24. Liang, M., Ye, H., Shen, Q., Jiang, X., Cui, G., Gu, W., Zhang, L.H., Naqvi, N.I. and Deng, Y.Z. (2021) Tangeretin inhibits fungal ferroptosis to suppress rice blast. J Integr Plant Biol, 63, 2136–2149.

25. Liu, C., Li, Z., Xing, J., Yang, J., Wang, Z., Zhang, H., Chen, D., Peng, Y.L. and Chen, X.L. (2018) Global analysis of sumoylation function reveals novel insights into development and appressorium-mediated infection of the rice blast fungus. New Phytol, 219, 1031–1047.

26. Liu, M., Wang, F., He, B., Hu, J., Dai, Y., Chen, W., Yi, M., Zhang, H., Ye, Y., Cui, Z., Zheng, X., Wang, P., Xing, W. and Zhang, Z. (2024) Targeting *Magnaporthe oryzae* effector MoErs1 and host papain-like protease OsRD21 interaction to combat rice blast. Nature plants. 10.1038/s41477-024-01642-x

27. Liu, X.H., Liang, S., Wei, Y.Y., Zhu, X.M., Li, L., Liu, P.P., Zheng, Q.X., Zhou, H.N., Zhang, Y., Mao, L.J., Fernandes, C.M., Del Poeta, M., Naqvi, N.I. and Lin, F.C. (2019) Metabolomics analysis identifies sphingolipids as key signaling moieties in appressorium morphogenesis and function in *Magnaporthe oryzae*. mBio, 10, e01467–01419.

28. Mamaev, D.V. and Zvyagilskaya, R.A. (2019) Mitophagy in yeast. Biochemistry, 84, S225–s232.

29. Mao, K., Wang, K., Zhao, M., Xu, T. and Klionsky, D.J. (2011) Two MAPK-signaling pathways are required for mitophagy in *Saccharomyces cerevisiae*. J. Cell Biol, 193, 755–767.

30. Meng, S., Jagernath, J.S., Luo, C., Shi, H. and Kou, Y. (2022) MoWhi2 mediates mitophagy to regulate conidiation and pathogenesis in *Magnaporthe oryzae*. Int. J. Mol. Sci, 23, 5311.

31. Oemer, G., Koch, J., Wohlfarter, Y., Lackner, K., Gebert, R.E.M., Geley, S., Zschocke, J. and Keller, M.A. (2022) The lipid environment modulates cardiolipin and phospholipid constitution in wild type and tafazzin-deficient cells. J. Inherit. Metab. Dis, 45, 38–50.

32. Osman, C., Haag, M., Wieland, F.T., Brügger, B. and Langer, T. (2010) A mitochondrial phosphatase required for cardiolipin biosynthesis: the PGP phosphatase Gep4. Embo j, 29, 1976–1987.

33. Park, Y., Cho, Y., Lee, Y.H., Lee, Y.W. and Rhee, S. (2016) Crystal structure and functional analysis of isocitrate lyases from *Magnaporthe oryzae* and *Fusarium graminearum*. J. Struct. Biol., 194, 395–403.

34. Rao, X., Huang, X., Zhou, Z. and Lin, X. (2013) An improvement of the 2^(-delta delta CT) method for quantitative real-time polymerase chain reaction data analysis. Biostat Bioinforma Biomath., 3, 71–85.

35. Rittenour, W.R., Chen, M., Cahoon, E.B. and Harris, S.D. (2011) Control of glucosylceramide production and morphogenesis by the Bar1 ceramide synthase in *Fusarium graminearum*. PloS one., 6, e19385.

36. Schäfer, F., Seip, N., Maertens, B., Block, H. and Kubicek, J. (2015) Purification of GST-Tagged Proteins. Methods in enzymology, 559, 127–139.

37. Schlame, M. and Greenberg, M.L. (2017) Biosynthesis, remodeling and turnover of mitochondrial cardiolipin. Biochim Biophys Acta Mol Cell Biol Lipids, 1862, 3–7.

38. Sentelle, R.D., Senkal, C.E., Jiang, W., Ponnusamy, S., Gencer, S., Selvam, S.P., Ramshesh, V.K., Peterson, Y.K., Lemasters, J.J., Szulc, Z.M., Bielawski, J. and Ogretmen, B. (2012) Ceramide targets autophagosomes to mitochondria and induces lethal mitophagy. Nat. Chem. Biol., 8, 831–838.

39. Sha, G., Sun, P., Kong, X., Han, X., Sun, Q., Fouillen, L., Zhao, J., Li, Y., Yang, L., Wang, Y., Gong, Q., Zhou, Y., Zhou, W., Jain, R., Gao, J., Huang, R., Chen, X., Zheng, L., Zhang, W., Qin, Z., Zhou, Q., Zeng, Q., Xie, K., Xu, J., Chiu, T.Y., Guo, L., Mortimer, J.C., Boutté, Y., Li, Q., Kang, Z., Ronald, P.C. and Li, G. (2023) Genome editing of a rice CDP-DAG synthase confers multipathogen resistance. Nature, 618, 1017–1023.

40. Shen, Z., Li, Y., Gasparski, A.N., Abeliovich, H. and Greenberg, M.L. (2017) Cardiolipin regulates mitophagy through the protein kinase C pathway. J. Biol. Chem., 292, 2916–2923.

41. Sweigard, J.A., Carroll, A.M., Farrall, L., Chumley, F.G. and Valent, B. (1998) *Magnaporthe grisea* pathogenicity genes obtained through insertional mutagenesis. Mol. Plant Microbe Interact., 11, 404–412.

42. Ting, H.C., Chen, L.T., Chen, J.Y., Huang, Y.L., Xin, R.C., Chan, J.F. and Hsu, Y.H. (2019) Double bonds of unsaturated fatty acids differentially regulate mitochondrial cardiolipin remodeling. Lipids Health Dis., 18, 53.

43. Wang, Z., Zhang, H., Liu, C., Xing, J. and Chen, X.L. (2018) A deubiquitinating enzyme Ubp14 is required for development, stress response, nutrient utilization, and pathogenesis of *Magnaporthe oryzae*. Front. Microbiol., 9, 769.

44. Wu, X.Y., Dong, B., Zhu, X.M., Cai, Y.Y., Li, L., Lu, J.P., Yu, B., Cheng, J.L., Xu, F., Bao, J.D., Wang, Y., Liu, X.H. and Lin, F.C. (2023) SP-141 targets Trs85 to inhibit rice blast fungus infection and functions as a potential broad-spectrum antifungal agent. Plant communications, 100724.

45. Wu, Y.X., Zhang, Y.D., Li, N., Wu, D.D., Li, Q.M., Chen, Y.Z., Zhang, G.C. and Yang, J. (2022) Inhibitory effect and mechanism of action of juniper essential oil on gray mold in cherry tomatoes. Front. Microbiol., 13, 1000526.

46. Xiao, Y., Liu, L., Zhang, T., Zhou, R., Ren, Y., Li, X., Shu, H., Ye, W., Zheng, X., Zhang, Z. and Zhang, H. (2021) Transcription factor MoMsn2 targets the putative 3-methylglutaconyl-CoA hydratase- encoding gene *MoAUH1* to govern infectious growth via mitochondrial fusion/fission balance in *Magnaporthe oryzae*. Environ Microbiol., 23, 774–790.

47. Xu, J.R., Staiger, C.J. and Hamer, J.E. (1998) Inactivation of the mitogen-activated protein kinase Mps1 from the rice blast fungus prevents penetration of host cells but allows activation of plant defense responses. Proc. Natl Acad Sci. USA., 95, 12713–12718.

48. Yamashita, S.I., Jin, X., Furukawa, K., Hamasaki, M., Nezu, A., Otera, H., Saigusa, T., Yoshimori, T., Sakai, Y., Mihara, K. and Kanki, T. (2016) Mitochondrial division occurs concurrently with autophagosome formation but independently of Drp1 during mitophagy. J. Cell Biol., 215, 649–665.

49. Yang, L., Zhao, M., Sha, G., Sun, Q., Gong, Q., Yang, Q., Xie, K., Yuan, M., Mortimer, J.C., Xie, W., Wei, T., Kang, Z. and Li, G. (2022) The genome of the rice variety LTH provides insight into its universal susceptibility mechanism to worldwide rice blast fungal strains. Comput. Struct. Biotechnol. J., 20, 1012–1026.

50. Yin, J., Hao, C., Niu, G., Wang, W., Wang, G., Xiang, P., Xu, J.R. and Zhang, X. (2020) FgPal1 regulates morphogenesis and pathogenesis in *Fusarium graminearum*. Environ Microbiol., 22, 5373–5386.

51. Yin, Z., Tang, W., Wang, J., Liu, X., Yang, L., Gao, C., Zhang, J., Zhang, H., Zheng, X., Wang, P. and Zhang, Z. (2016) Phosphodiesterase MoPdeH targets MoMck1 of the conserved mitogen-activated protein (MAP) kinase signalling pathway to regulate cell wall integrity in rice blast fungus *Magnaporthe oryzae*. Mol. Plant Pathol., 17, 654–668.

52. Yuan, H., Yuan, M., Shi, B., Wang, Z., Huang, T., Qin, G., Hou, H., Wang, L. and Tu, H. (2022) Biocontrol activity and action mechanism of *Paenibacillus polymyxa* strain Nl4 against pear Valsa canker caused by *Valsa pyri*. Front. Microbiol., 13, 950742.

53. Zheng, W., Zhou, J., He, Y., Xie, Q., Chen, A., Zheng, H., Shi, L., Zhao, X., Zhang, C., Huang, Q., Fang, K., Lu, G., Ebbole, D.J., Li, G., Naqvi, N.I. and Wang, Z. (2015) Retromer is essential for autophagy-dependent plant infection by the rice blast fungus. PLoS Genet., 11, e1005704.

