## Supplemental File for "Inhibitor of cardiolipin biosynthesis-related enzyme MoGep4 confers broad-spectrum anti-fungal activity"

The following Supporting Information is available for this article:

**Figure S1.** Cardiolipin (CL) biosynthesis pathway and generation of mutants for genes *MoGEP4*, *MoCLD1* and *MoTAZ1* in *M. oryzae*.

**Figure S2** *MoGEP4* is required for aerial hyphae growth, and stress responses in *M. oryzae*.

**Figure S3.** MoGep4 is required for lipid and glycogen translocation and degradation.

**Figure S4.** Subcellular localization of MoGep4.

**Figure S5.** Expression of *MoGEP4*$^{D60A}$ does not rescue the defect of the *Mogep4* mutant.

**Figure S6.** Gep4 protein structures of *M. oryzae* and other pathogens.

**Figure S7.** Alexidine dihydrochloride has no side effects on wheat and rice seedlings.

**Table S1.** Wild-type and mutant strains of *Magnaporthe oryzae* used in this study.

**Table S2.** Primers used in this study.

**Supporting Information**

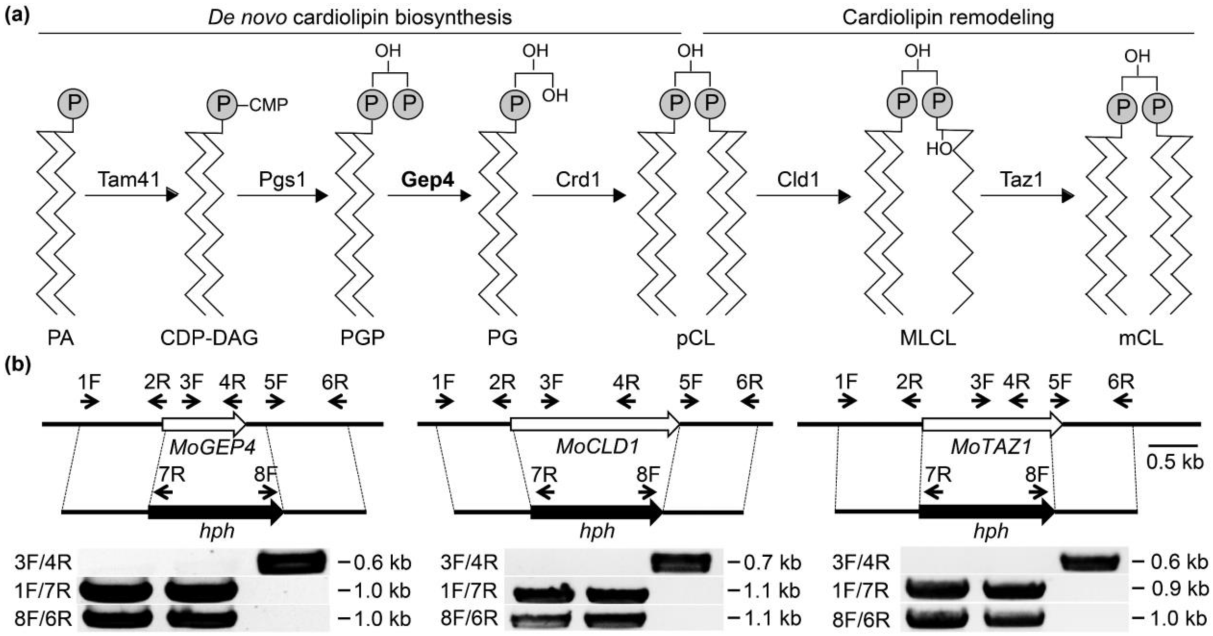


**Figure S1. Cardiolipin (CL) biosynthesis pathway and generation of mutants for genes *MoGEP4, MoCLD1* and *MoTAZ1* in *M. oryzae*.**

**(a)** *de novo* CL biosynthesis and CL remodeling pathway, which is adapted from yeast. PA, phosphatidic acid; CDP-DAG, CDP-diacylglycerol; PGP, phosphatidylglycerophosphate; PG, phosphatidylglycerol; pCL, precursor cardiolipin; MLCL, monolyso-cardiolipin; mCL, mature cardiolipin. **(b)** Primer pairs 1F/7R and 8F/6R were used to amplify upstream and downstream flanking fragments for gene deletion, and primer pair 3F/4R was used to screen for the knockout mutant.

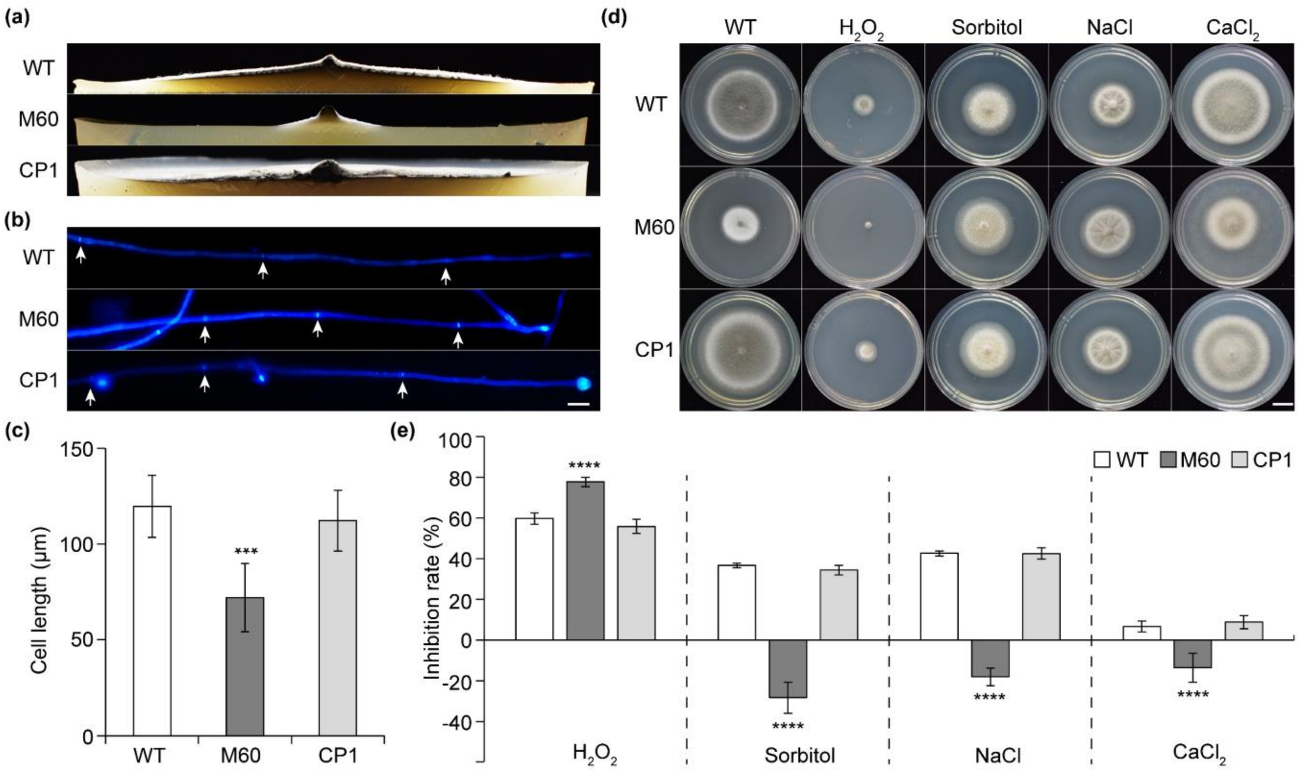


**Figure S2. *MoGEP4* is required for aerial hyphae growth, and stress responses in *M. oryzae*.**

**(a)** Radial growth of colonies on CM plates at 5 dpi. **(b)** Hyphal tips of the WT, M60, and CP1 strains were stained with Calcofluor White (CFW). Cell septa are indicated by white arrows. Bar, 20 µm. **(c)** Apical and subapical cell length of the WT, M60, and CP1 strains. Bar, 10 µm. Five-day-old colony **(d)** and inhibition rates **(e)** of the WT, M60, and CP1 strains on CM supplemented with different stressors, including $H_2O_2$ (0.01 M), sorbitol (1 M), NaCl (0.7 M), and $CaCl_2$ (0.1 M). Bar, 1 cm. Error bars in **(c, e)** indicate standard deviations, and asterisks indicate statistically significant differences using the unpaired Student's *t*-test (***$P < 0.001$; ****$P < 0.0001$).

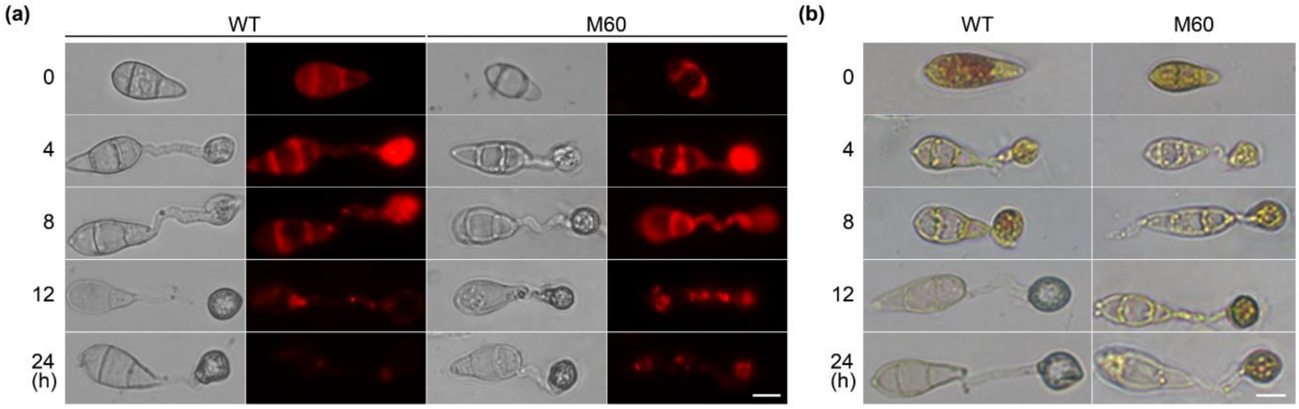


**Figure S3. MoGep4 is required for lipid and glycogen translocation and degradation.**

**(a)** Microscopic visualization of lipid droplets during appressorium formation. Nile Red solution was used for lipid staining. Bar, 10 µm. **(b)** The WT and *Mogep4* (M60) strain were stained with the iodine solution at different time points. Glycogen was visualized as yellowish-brown deposits. Bar, 10 µm.

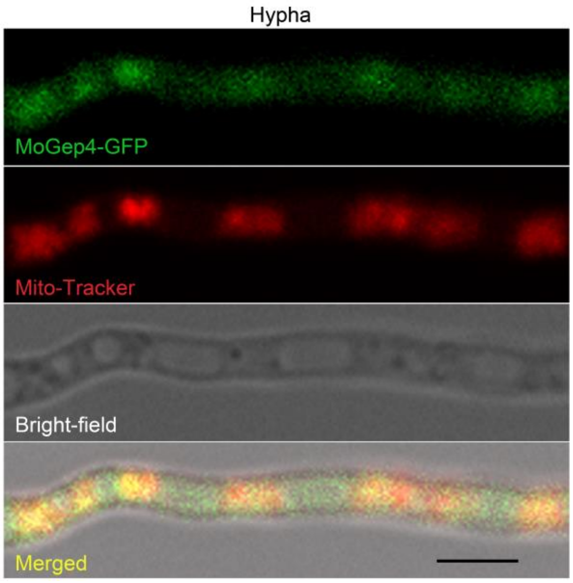


**Figure S4. Subcellular localization of MoGep4.**

MoGep4 localizes to mitochondria in hyphae. The strains were cultured in the CM for confocal observation at 36 h. Mitochondrial dye was used for Mito-Tracker. Bars, 5 μm.

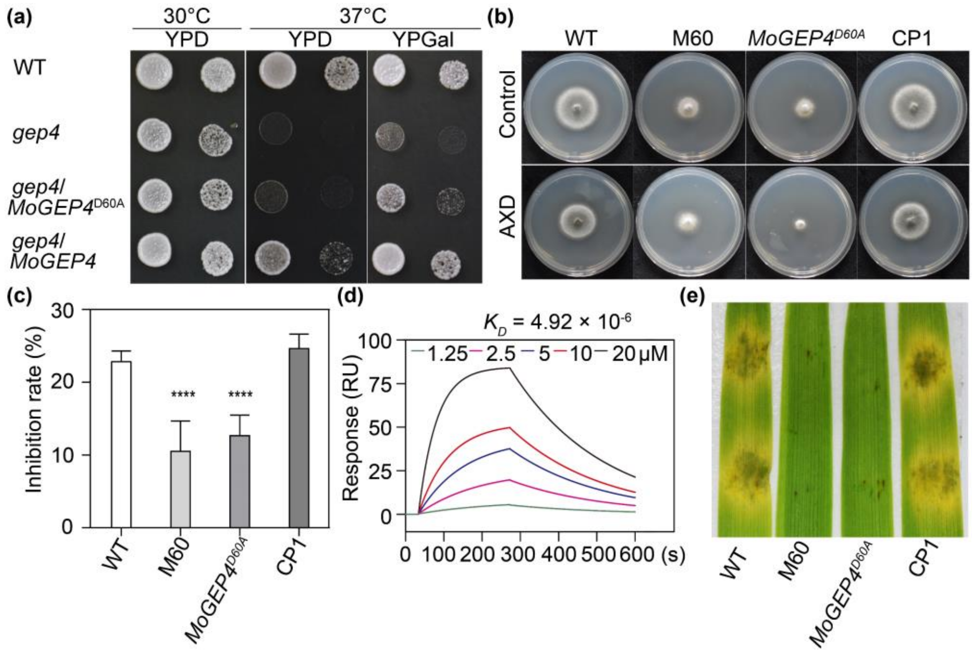


**Figure S5. Expression of *MoGEP4$^{D60A}$* does not rescue the defect of the *Mogep4* mutant.**

**(a)** Yeast strains BY4741 (WT), the *gep4* mutant (*gep4*), transformants of the *gep4* mutant carrying pYES2-*MoGEP4* (*gep4*/*MoGEP4*) or pYES2-*MoGEP4*$^{D60A}$ (*gep4*/*MoGEP4$^{D60A}$*) were cultured on YPD and YPGal plates at 30°C and 37°C for 4 days, respectively. **(b)** Five-day-old cultures of the wild-type (WT), *Mogep4* mutant (M60), *GEP4$^{D60A}$*(*gep4*/*MoGEP4$^{D60A}$*), and *Mogep4* complementation (CP1) strains on CM plates, supplemented with/without AXD (4 μM). **(c)** Inhibition rates of AXD shown in **(b)**. **(d)** SPR analysis of the binding affinity of AXD to MoGep4$^{D60A}$-GST and the GST tag as the control shown in **(Figure 4c)**. **(e)** Drop-inoculation of the WT, M60, *GEP4$^{D60A}$*, and CP1 strains on barley leaves at 5 days post-inoculation (dpi). Means and standard deviations were calculated from three replicates, and asterisks indicate statistically significant differences using the Student's *t*-test (****$P$ < 0.0001).

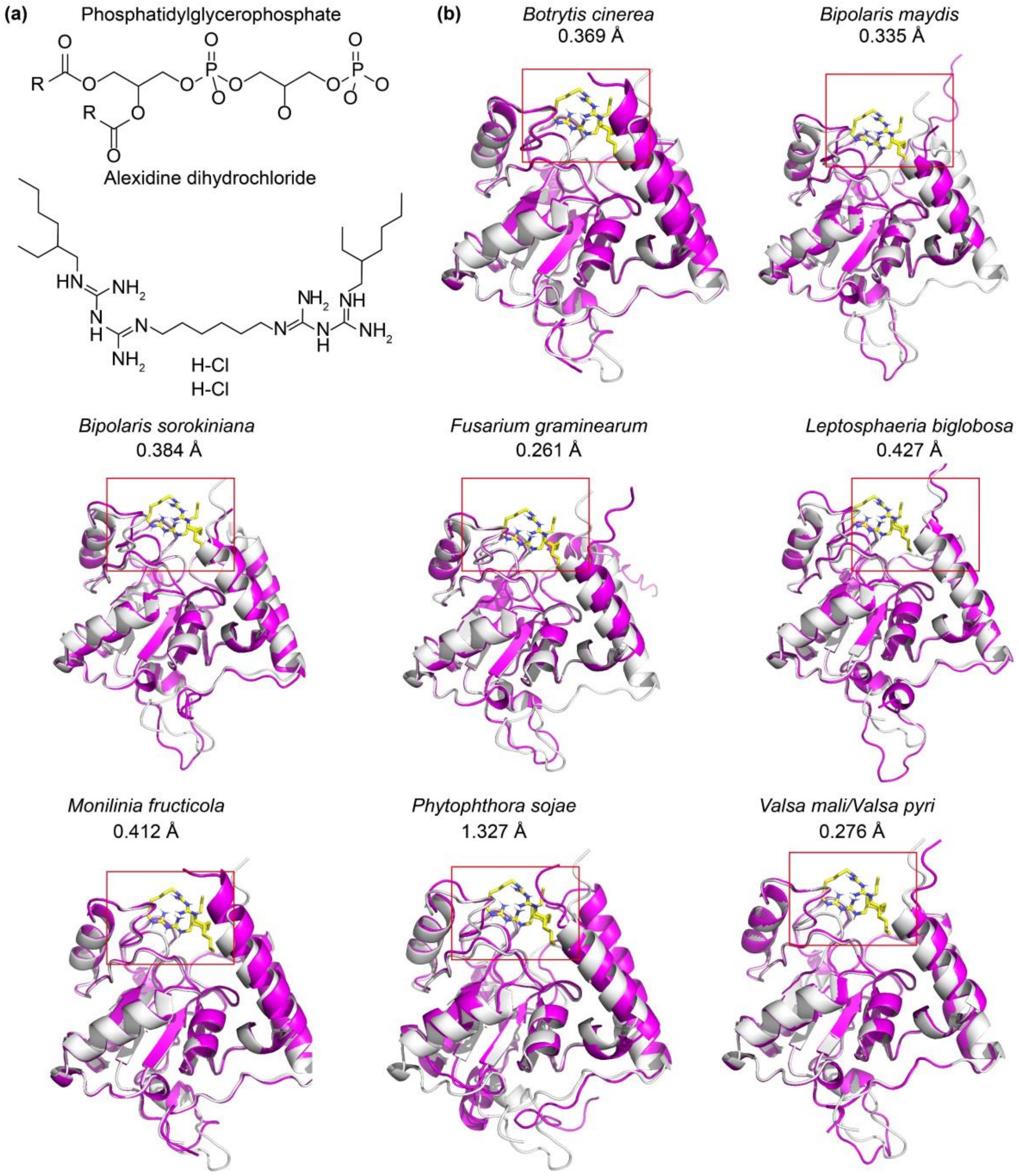


**Figure S6. Gep4 protein structures of *M. oryzae* and other pathogens.**

**(a)** Chemical structures of phosphatidylglycerophosphate and alexidine dihydrochloride. **(b)** Gray indicates the predicted protein structure of MoGep4 and green for predicted structures of Gep4 homologs. The binding pocket is highlighted by the red box. The number denotes root mean square deviation (RMSD).

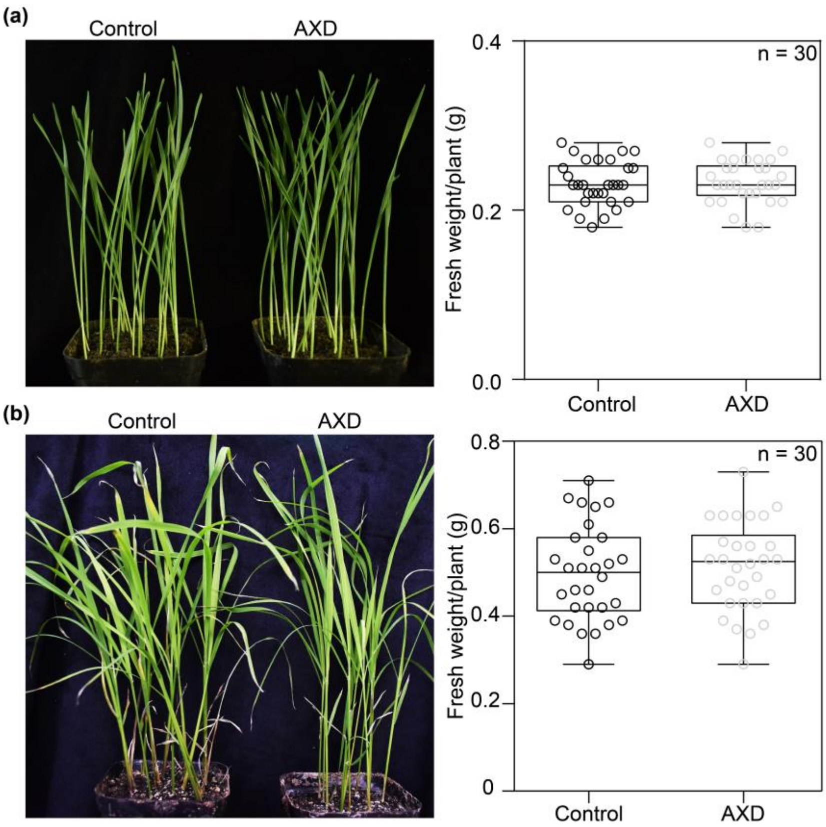


**Figure S7. Alexidine dihydrochloride has no side effects on wheat and rice seedlings.**

**(a)** Seven-day-old wheat seedlings exposed to alexidine dihydrochloride (AXD, 4 μM) for three days and fresh weight per plant. **(b)** Four-week-old rice seedlings exposed to AXD (4 μM) for three days and fresh weight per plant. Solvent dimethyl sulfoxide (DMSO) was used as the control.

**Table S1. Wild-type and mutant strains of *Magnaporthe oryzae* used in this study.**

| strain | Genotype description | Reference |
|---|---|---|
| P131 | Wild type strain | This study |
| M60 | *MoGEP4* deletion mutant of *M. oryzae* | This study |
| *Moatg24* | *MoATG24*deletion mutant of *M. oryzae* | This study |
| *cld1* | *MoCLD1* deletion mutant of *M. oryzae* | This study |
| *taz1* | *MoTAZ1* deletion mutant of *M. oryzae* | This study |
| CP1 | *Mogep4* mutant expression *MoGEP4*-GFP | This study |
| WT/MoIdh1-GFP | WT expression *MoIDH1*-GFP | This study |
| M60/MoIdh1-GFP | *Mogep4* mutant expression *MoIDH1*-GFP | This study |
| *ATG24*-GFP | *Moatg24* expression *MoATG24*-GFP | This study |
| BPA1 | WT expression *MoATG24*-nYFP and *PKC*-cYFP | This study |
| *px* | *Moatg24* expression *MoATG24* $^{\Delta PX}$ | This study |
| *bar* | *Moatg24* expression *MoATG24* $^{\Delta BAR}$ | This study |
| WT/*ATG24*-GFP+*PKC1*-3×FLAG | WT expression *MoATG24*-GFP and *PKC*-3×FLAG | This study |
| M60/*ATG24*-GFP+*PKC1*-3×FLAG | *Mogep4* mutant expression *MoATG24*-GFP and *PKC*-3×FLAG | This study |
| GOX1 | Overexpression of *MoGEP4* in *M. oryzae* | This study |

**Table S2. Primers used in this study.**

| Name | Sequence (5' > 3') |
|---|---|
| AD- PKC1/R | tgcagctcgagctcgatggatcctcaatcaaagtctgccgtgtac |
| AD-Mkk1/F | acgtaccagattacgctcatatgatgcacgatcaagaagctgcc |
| AD-Mkk1/R | tgcagctcgagctcgatggatcctcactctgcaggcttggct |
| AD-PKC1/F | acgtaccagattacgctcatatgatggatgacaggatacaagacatt |
| ATG24-1F | ctgtttgcctggtgaagtgagt |
| ATG24-2R | ttgacctccactagctccagccaagccgctggtgtagttaggctctggt |
| ATG24-3F | gaatagagtagatgccgaccgcgggttcacggacttttacgacgcca |
| ATG24-4R | tgtattgatgcggtttcttgct |
| ATG24-5F | gccgactttcaaaaacccga |
| ATG24-6R | ctcgcaggcttccaagataa |
| ATG24-7F | gttttaggcaacggcacgac |
| ATG24-8R | cggcttgagagattgacgag |
| ATG24BAR-BDR1 | cggccgctgcaggtcgacggatccctaagcagccacggcgccctcggcgttgatgaaggtatcgg |
| ATG24-GFP-F | agggaacaaaagctgggtaccacccaaaccgaccagtaagc |
| ATG24-GFP-R | gcccttgctcaccataagcttagcagccacggcgccctc |
| ATG24-NYFP-F | agggaacaaaagctgggtaccgccctcctttcaacaaccag |
| ATG24-NYFP-R | cgtggcgatggagcgaagcttagcagccacggcgccctc |
| ATG24PX-BDF | tctcagaggaggacctgcatatgatggggggaatcgaccaaga |
| ATG24PX-BDF1 | ctagtaactacacatgactggaacgccacgatgag |
| ATG24PX-BDR1 | cgtggcgttccagtcgcccaggcccatgccgtg |
| BD- ATG24/F | tctcagaggaggacctgcatatgatggggggaatcgaccaaga |
| BD- ATG24/R | cggccgctgcaggtcgacggatccctaagcagccacggcgc |
| CLD1-1F | gcacccagtgttagatttacgag |
| CLD1-2R | ttgacctccactagctccagccaagccccgctggcagtcgaagaagt |
| CLD1-3F | cgcacaatgtcggcttactt |
| CLD1-4R | cgccactcctcaagagcatc |
| CLD1-5F | gaatagagtagatgccgaccgcgggttttcgcaaggagaaatcggca |
| CLD1-6R | gacgacgagccccaggatta |

| | |
|---|---|
| GEP-2R | ttgacctccactagctccagccaagccgcgatgctgaaaggttgagg |
| GEP4/2KOF2 | cgattcgatactaacgccgccaaggccggaaactaaccc |
| GEP4/2KOR2 | ggccctattcatccgtga |
| GEP4/KO1F3 | atgagagacttcgcgcca |
| GEP4/KOF1 | gccatcccagcaagtatctt |
| GEP4/KOF1 | gccatcccagcaagtatctt |
| GEP4/KOF2 | cgattcgatactaacgccgccaaggccggaaactaaccc |
| GEP4/KOR1 | ccatgtcgctggccgggtgacactgccgcctttcagttt |
| GEP4/KOR1 | ccatgtcgctggccgggtgacactgccgcctttcagttt |
| GEP4/KOR2 | aatccacagttacgtgagatgg |
| GEP4-1F | cgctctgtgacttggacggt |
| GEP4-3F | acaaatgtcatccacgagcc |
| GEP4-4R | cttctgaggaggaacggcat |
| GEP4-5F | gaatagagtagatgccgaccgcgggttggcaggctaaatactccacg |
| GEP4-6R | cactgggctccaaccattct |
| GEP4-Com-F | agggaacaaaagctgggtaccccacagcagtgagaaatcgg |
| GEP4-D60A-F | gttattctggataaggcagactgtttcgcatacccaga |
| GEP4-D60A-R | atgcgaaacagtctgccttatccagaataactgccttt |
| GEP4-GFP-R | gcccttgctcaccataagcttttcaaaaggactttgtggttgg |
| GEP4KO2/R4 | aatccacagttacgtgagatgg |
| GEP4-RP27-F | aacccaatcttcaaactcgagatgaacctcaacctttcagca |
| GEP4-RP27-R | gcccttgctcaccataagcttacggggcgtggagtatttag |
| GEP4SCF1 | tcttcctataccaatccacgat |
| GEP4SCF1 | tcttcctataccaatccacgat |
| GEP4SCR1 | tgccgaaactttaacaccat |
| GEP4SCR1 | tgccgaaactttaacaccat |
| H850 | ttgtccgtcaggacattgtt |
| H852 | aactcaccgcgacgtctgtc |
| H855R: | gctgatctgaccagttgc |
| H856F | gtcgatgcgacgcaatcgt |

| | |
|---|---|
| HY/R: | gtattgaccgattccttgcggtccgaa |
| HYG/F | ggcttggctggagctagtggaggtcaa |
| HYG/R | aacccgcggtcggcatctactctattc |
| IDH-GFP-F | agggaacaaaagctgggtaccgctgcccctaatgactgtaaac |
| IDH-GFP-R | gcccttgctcaccataagcttgaaagtctccatcttgtccaag |
| KanMXF | gtcacccggccagcgacatgg |
| KanMXR | ggcggcgttagtatcgaatcg |
| MGG02252_QF | acgccgtctactcaggatca |
| MGG02252_QR | tctcgccgtttggaatgtat |
| MGG05059_QF | cgactccaaggactgggata |
| MGG05059_QR | gtcctcggacaccttctcc |
| MGG07219_QF | gcaatgtcggtcccaactac |
| MGG07219_QR | atctcaaaggcgatgacacc |
| PKC1-3xFLAG-F | aaacccgggctgcaggaattcaggtctgctgaggtctgtctta |
| PKC1-3xFLAG-R | gtcgacggtatcgataagcttatcaaagtctgccgtgtacga |
| PKC1-CYFP-F | agggaacaaaagctgggtaccacgcacagtgacacaaagattcg |
| PKC1-CYFP-R | cttgcaggccgggcgaagcttatcaaagtctgccgtgtacg |
| TAZ1-1F | tcctgtcagatggtgtgcga |
| TAZ1-2R | ttgacctccactagctccagccaagccgccgtcacccccatttttgt |
| TAZ1-3F | gcccacgacatttgctttca |
| TAZ1-4R | agggcgtcccactttcgttt |
| TAZ1-5F | gaatagagtagatgccgaccgcgggttgttctcaaacttcggcacgg |
| TAZ1-6R | cttacaacggagcacaggga |
| TubQF | tctgacttcaggaatggtcgttac |
| TubQR | agcggtctggatgttgttgg |
| YG/F | gatgtaggagggcgtggatatgtcct |
